# The SynMuvA *lin-15A* licenses natural transdifferentiation by antagonizing identity safeguarding mechanisms

**DOI:** 10.64898/2026.01.22.701184

**Authors:** Sarah Becker, Marie-Charlotte Morin, Julien Lambert, Shashi Kumar Suman, Francesco Carelli, Alex Appert, Stéphane Roth, Sarah Hoff-Yoessle, Jessica Medina-Sanchez, Manuela Portoso, Stéphanie Le Gras, Julie Ahringer, Sophie Jarriault

## Abstract

The mechanisms that restrict or enable latent cellular plasticity have attracted growing interest over the past decade, with important implications for cancer and regenerative therapies. However, the diversity of both pro- and anti-plasticity mechanisms remains incompletely understood. Here, we identify the THAP domain gene *lin-15A* as a novel factor involved in the natural rectal-to-neuronal Y-to-PDA transdifferentiation in *Caenorhabditis elegans*. We found that, unlike previously described essential factors, *lin-15A* is not a Driver of transdifferentiation. Instead, it antagonizes several chromatin-modifying complexes known to safeguard differentiated cell identities. We also show that *lin-15A* is not a core plasticity factor per se but acts as one specifically in the Y cell context. Together, our findings support a model in which diverse molecular players coordinate controlled cell identity conversions: plasticity factors function as Drivers, while others like *lin-15A* which we propose to term Licensers attenuate identity safeguarding mechanisms, thereby facilitating transdifferentiation.

## Introduction

Over 60 years ago, John Gurdon demonstrated that nuclei of differentiated, somatic cells can be reprogrammed and allow the development of a frog, when inserted into an enucleated frog egg (Gurdon, 1960, 1962). Since then, numerous studies have shown that transcription factor-based inducing cues, ranging from only one to a small cocktail of TFs, could trigger the reprogramming of various differentiated cells into ES-like cells, as well as into another differentiated identity (aka induced direct reprogramming or induced transdifferentiation), both *in vitro* (Takahashi & Yamanaka, 2016; Wang et al., 2021) and *in vivo* (Zhang et al., 2022; Rothman & Jarriault, 2019). The control and plasticity of differentiated cell identities have emerged as central topics in biomedical research, particularly in the contexts of cancer, tissue regeneration, and cellular reprogramming (Yuan et al., 2019; Takahashi and Yamanaka, 2006). Understanding how fully differentiated cells can shift their identity, and why not all cells manage to do so, offers valuable insights into developmental biology, disease progression, and therapeutic innovation.

In the past few decades, extensive research using induced reprogramming systems has highlighted several roadblocks. For instance, induced direct reprogramming typically exhibit low efficiency, yielding only a small fraction of successfully converted cells, and the reprogramming efficiency varies widely between cell types, suggesting the presence of intrinsic barriers. In addition, induced direct reprogramming is more easily achieved by over-expressing specific transcription factors in cells that are not fully differentiated compared to those that are differentiated (Wang et al., 2021; Rothman and Jarriault, 2019). Moreover, conversion is often incomplete, with cells retaining an epigenetic memory or original lineage gene expression, or yields immature cells. These observations have led to the concept of reprogramming barriers.

A number of mechanisms have been proposed to explain the resistance of most differentiated cells to induced reprogramming. Extra-cellular matrix properties (Huels & Medema, 2018), cell signaling pathways (Apostolou & Hochedlinger; 2013), cell identity acquisition and maintenance transcription factors (TFs) (Patel & Hobert, 2017), fate-stabilizing TFs (Mall et al., 2017; Lee et al., 2019; Missinato et al., 2023), as well as genome organization, including 3D chromatin architecture, histone modifications and chromatin remodelers (Wang et al., 2021; Özcan & Tursun, 2023), have all been proposed to modulate the response of differentiated cells to reprogramming cues. These mechanisms may altogether underlie the active safeguarding of the differentiated identities suggested by (Blau and Baltimore, 1991).

Do barriers exist in cells that naturally undergo direct reprogramming? What would these barriers be and how are they removed? Natural instances of direct reprogramming, in which a differentiated cell acquires itself or gives rise to a daughter cell with a different differentiated identity without a passage through a stable intermediate, are not very frequent in a given species. In *C. elegans*, although some cells naturally change their identity during development, the majority retain their identity. In other model systems as well, including vertebrates, most cell do not naturally change their identity apart from some specific developmental, regenerative or pathological contexts. However, by contrast with induced direct reprogramming, natural direct reprogramming (aka natural transdifferentiation, or Td), is robust and yields a stable and mature final cell (Zuryn et al., 2014; Sousounis et al., 2015). Thus, natural Td presents a compelling model of endogenous cell fate flexibility operating outside of artificial manipulation (Eguchi et al., 1974; Selman and Kafatos, 1974; Okada, 1991; Henry and Tsonis, 2010; Kodama and Eguchi, 1995; Yamada, 1977; Schmid and Reber-Müller, 1995; Merrell and Stanger, 2016).

Here, we have used an instance of natural Td in *Caenorhabditis elegans* to address these questions. Given its transparency, complete toolbox for genetical approaches and fully resolved stereotypical somatic lineages (Sulston and Horvitz, 1977) it is possible to unambiguously examine cellular identities at the single cell level in *C. elegans*, making it a prime model for the study of natural Td (Jarriault et al., 2008; Riddle et al., 2013; Sammut et al., 2015; Molina-Garcia et al., 2020; Riva et al., 2022, Rothman and Jarriault, 2019). The transdifferentiation of the rectal Y cell into the cholinergic motorneuron PDA in the hermaphrodite was the first evidence of natural Td in the worm and is the best characterized event to this day (Jarriault et al., 2008; reviewed in Lambert et al., 2021) (Fig 1A). This specialized epithelial cell is born during embryogenesis and forms the rectal tube together with 5 other rectal cells: B, U, F, K and K’ that share similar morphology and markers (Sulston and Horvitz, 1977). At the end of the L1 larval stage however, the rectal Y retracts from the rectum (Td initiation), dedifferentiates very transiently into an unipotent intermediate, and re-differentiates into the PDA neuron in a stereotypical and robust process (Richard et al., 2011; Zuryn et al., 2014). Y-to-PDA occurs directly and without a cell division (Jarriault et al., 2008; White et al., 1986). The erasure of the initial rectal identity requires the action of conserved NODE (Nanog and Oct4-associated deacetylase)-like complex (Kagias et al., 2012) while the H3K4me3 histone methyltransferase SET-2/SET1 ensures Td robustness in face of environmental stresses (Zuryn et al., 2014). The NODE-like complex, composed of SEM-4/SALL4, CEH-6/OCT, EGL-27/MTA1 acts together with the region-specific HOX TF EGL-5, the conserved plasticity factor SOX-2/SOX2 and the bHLH transcription factor HLH-16 (Daniele et al., 2025; Rashid et al., 2022), as the Drivers of the initiation step (Kagias et al., 2012). The loss of any of these factors result in the failure of Td initiation: no retraction of Y from the rectum and maintenance of the initial rectal identity. ZTF-11/Myt1 is also required for this step, however its relation to the Drivers is still unknown (Lee et al., 2019). Re-differentiation into PDA is dependent on UNC-3/EBF (Richard et al., 2011) acting as a terminal selector, and facilitated by the histone demethylase JMJD-3.1 (Zuryn et al., 2014).

**Figure 1:**
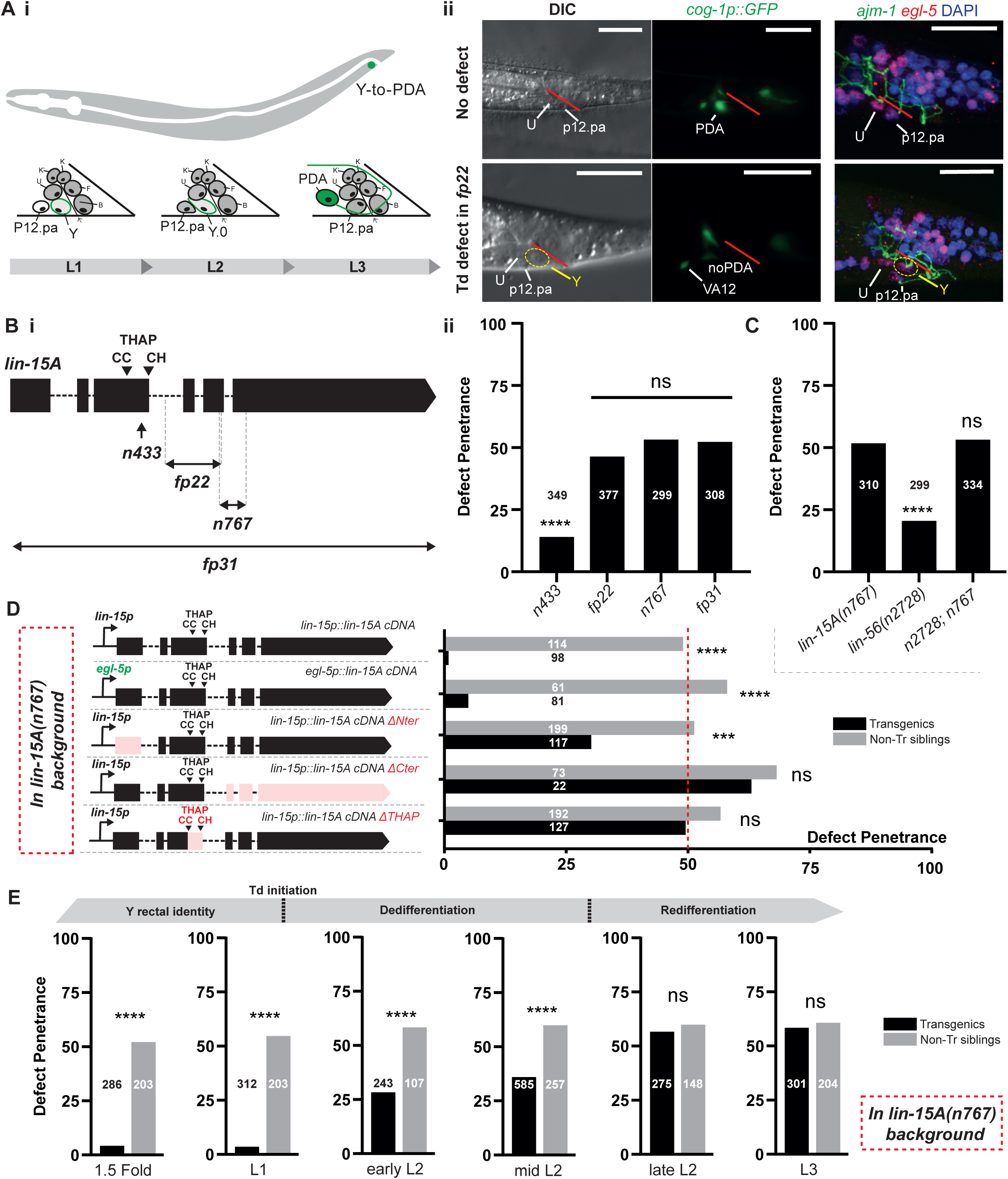
LIN-15A and LIN-56 are two novel factors required for Y-to-PDA initiation. **A i** Y-to-PDA transdifferentiation and typical defects of *fp22* mutants. At the end of L1 stage, Y retracts from the rectum and dedifferentiates. PDA redifferentiation takes place during the L2 and late L3 stages. **ii** Defects induced by fp22 result in the inability of the Y cell to retract from the rectum and intiate its Td - 3 epithelial nuclei visible in DIC and initiate Y-to-PDA. Images of the rectal area, the rectal slit is indicated in red. iii In Y-to-PDA defective fp22 animals, the Y cell retains its epithelial features (tight junctions’ component AJM-1) and markers (LIN-26). Scalebars = 10 µm. **B i** Alleles of *lin-15A* used in this study and their respective defect penetrance. **ii** *fp22*, *fp31 and n767* are null alleles and induce a 50% defect penetrance in Td initiation. Weaker *433* causes defects at a penetrance of 15%. X2 and Fisher exact tests with Bonferroni corrections for proportion comparisons. For this entire study, the numbers associated with the histograms or scatter plots indicate the relative n of the cohort. **C** *lin-15A* and *lin-56* act in cooperation for Y-to-PDA Td. X2 and Fisher exact tests with Bonferroni corrections for proportion comparisons. **D** Rectal-specific rescue of *lin-15A* by *egl-5* promoter fully suppresses the transdifferentiation defects demonstrating that *lin-15A* activity in the rectal cells is sufficient for its role in Y-to-PDA Td. The C-term and THAP domain of LIN-15A are critical for its role in Y-to-PDA Td. Its N-term domain is less critical. Experiment carried out in *lin-15A(n767)* background, with the standard 50% defect penetrance of these mutants indicated by the vertical red dotted line. Fisher exact tests with Bonferroni corrections for proportion comparisons. **E** Timing of LIN-15A requirement in Y-to-PDA transdifferentiation using a heat-shock inducible promoter. LIN-15A activity is only required during the L1 stage. LIN-15A rescue during the dedifferentiation step (L2 stage) partially supress the defects highlighting a possible timeframe of initiation spanning until the mid-L2 stage. LIN-15A rescue after the initiation step has no effect (late L2 and L3 stages). Experiment carried out in *lin-15A(n767)* background. Fisher exact tests with Bonferroni corrections for proportion comparisons.

In this study, we identify the SynMuvAs LIN-15A and LIN-56 as novel players in Y-to-PDA initiation, acting cooperatively. *lin-15A* is part of the *lin-15* operon together with the SynMuvB gene *lin-15B* (Clark et al., 1994; Huang et al., 1994) and codes for a THAP domain ZnF nuclear protein whose function is unknown (Davison et al., 2011). We report that it binds naked and nucleosome-bound chromatin, that its action in Td occurs in the rectal cells and requires its THAP domain. Our genetic interaction studies demonstrate that LIN-15A and LIN-56 function in a parallel pathway to the factors of the NODE-like complex. We also show that LIN-15A and LIN-56 are similarly required for Y-to-PDA in males (which occurs through Y division), but that contrary to the NODE-like complex, LIN-15A is not involved in the other rectal-to-neuronal transdifferentiation K-to-DVB (Riva et al., 2022), and that it can even restrict cellular plasticity in the embryo. Hence, LIN-15A acts as a plasticity factor in specific cell contexts, like that of the Y cell, whether in presence (males) or in absence (hermaphrodites) of a cell division. In contrast to VPC specification, we show that the role of LIN-15A is independent of *lin-3/EGF* during Td. However, we found that it antagonizes most of the SynMuvBs chromatin factors and complexes, like the DREAM complex, that negatively regulate chromatin activity, suggesting a role in the control of gene expression. SynMuvB factors appear to act as brakes for Y-to-PDA transdifferentiation, likely by safeguarding the initial rectal identity. Last, our ChIP-seq data suggest that the antagonism between LIN-15A and DREAM occurs at the chromatin level, with LIN-15A restricting DREAM binding on a subset of target genes.

## Results

### Two novel factors required for Y-to-PDA transdifferentiation: LIN-15A and LIN-56

Via an unbiased EMS screen for mutants inducing defects in Y-to-PDA transdifferentiation (Td) (PDA neuron not produced), we retrieved the allele *fp22* which leads to approximatively 50% of the animals failing to produce PDA (Fig 1.Aii and 1.B). EMS-density mapping using WGS analysis (Zuryn et al., 2010) identified *fp22* to be a 502 bp deletion in the coding sequence of *lin-15A* (Fig 1.B, See Appendix). The two canonical alleles of *lin-15A*, *n433* (weak loss of function) and *n767* (a 178 bp deletion-insertion in the C-term domain of LIN-15A, supposed null (Davison et al., 2011) equally result in Y-to-PDA defects with a penetrance of respectively 15% and 50% (Fig 1.B), confirming that *fp22* is a novel *lin-15A* allele (Fig 1.B and 1.D). Although *n767* behaves as a null allele, the molecular lesion does not entirely eliminate *lin-15A* but results in a truncated protein, leaving the possibility that the 50% defect penetrance observed could be due to residual *lin-15A* activity. We thus generated *fp31*: a deletion of the entire *lin-15A* gene (keeping *lin-15B* unaffected). As for *fp22* and *n767*, *fp31* resulted in a 50% penetrant Y-to-PDA defects (Fig 1.B). Genetic complementation tests further validated that *fp22* and *n767* are null alleles of *lin-15A* (Fig S1.A). For the rest of this study, experiments were conducted with the canonical *n767* allele unless otherwise stated.

LIN-15A was previously described to cooperate with LIN-56 (Davison et al 2011) in the repression of *lin-3/EGF* expression during VPC specification (Clark et al., 1994; Huang et al., 1994). Therefore, we tested whether a similar interaction occurs during Y-to-PDA. We found a 20% penetrance of Y-to-PDA defects in *lin-56(n2728)* null mutants (Davison et al., 2011). Double mutants *lin-56(n2728); lin-15A(n767)* displayed the same defect penetrance of 50% than single *lin-15A(n767)* mutants (Fig 1.C) indicating that the two factors act in the same pathway during Y-to-PDA. Our western blot experiments suggest that the LIN-15A-LIN-56 dimer is very stable (Figure S1.B).

Other SynMuvA genes, in addition to LIN-15A and LIN-56, repress the expression of *lin-3/EGF* during VPC specification (Fay and Yochem, 2007). Thus, we tested the putative involvement of other SynMuvAs in Y-to-PDA. However, none of the other tested SynMuvA appears to be required for Y-to-PDA (Fig S1.C).

Together, these results demonstrate the requirement of two novel factors in Y-to-PDA transdifferentiation: LIN-15A and LIN-56, where, contrary to VPC specification, they act independently of other SynMuvAs.

### LIN-15A acts through its THAP-domain, at the time of Y-to-PDA initiation, in the rectal cells

A defect in Y-to-PDA could result by a failure in the initiation step, where Y is retracted from the rectum, dedifferentiates and loses its epithelial features, or the re-differentiation step during which it becomes PDA and acquires neuronal features (Jarriault et al., 2008; Richard et al., 2011). DIC images of *lin-15A(fp22)* L3 and older worms (when Td should have taken place already) showed that the Y cell is retained in the rectum with apparent rectal morphology, a class I defect (Richard et al., 2011), indicating a Td initiation defect (Fig 1.Aii). Immunostainings against the epithelial and rectal markers, AJM-1 and EGL-5, respectively, confirmed that Y still carries rectal epithelial features and remains on the anterior side of the rectum, beyond the time point of Td initiation in Y-to-PDA defective worms (9/9 Y-to-PDA defective out of 21 animals scored) (Fig 1.Ai). Live reporters for additional rectal-epithelial markers showed that the persistent Y cell retains a rectal identity and that expression of these rectal markers was unaltered prior to Td initiation (Fig S1.D). Hence, LIN-15A is required for the initiation step. Of note, LIN-15A is expressed in Y prior to Td initiation (Fig S1.F), consistent with a role in promoting the identity switch.

Since *lin-15A* is broadly expressed, including in the rectal cells, during the earlier stages of *C. elegans* development (Fig S1.F), we sought to assess in which tissues and at what stage it was required for Td. Expression of *lin-15A* in the rectal cells using the *egl-5p* promoter was able to rescue the transdifferentiation defects to the same extent as when the *lin-15p* promoter is used as a driver (Fig 1.D and S1.E), indicating that LIN-15A acts the rectal cells during Y-to-PDA. We further found that both the C-terminal domain as well as the C2CH THAP domain are necessary for the function of LIN-15A for Td (Fig 1.D and S1.F). In contrast, we found that the N-terminal domain was only partially required since expressing a N-terminal truncated *lin-15A* cDNA could only partially rescue the defects. The requirement of the THAP domain suggests a nucleic acid binding role for LIN-15A in Y-to-PDA. In accordance with this hypothesis, CRISPR/Cas9 mCherry-tagged LIN-15A localizes in the nucleus, and purified LIN-15A is able to bind naked and nucleosome loaded DNA (Fig 1.F and S6).

We then assessed when LIN-15A activity was required for Y-to-PDA initiation. Y is born around 300 minutes after fertilization (during embryogenesis; Sulston et al., 1983), and Y-to-PDA initiation only occurs at the end of the first larval stage (Jarriault et al., 2008). Hence LIN-15A activity could be required at any time during that period as it is constantly expressed in the rectal cells until the transdifferentiation (Fig S1.E). Using a heat-shock (hs) inducible *lin-15A cDNA* transgene, we found that LIN-15A activity is required around Y-to-PDA Td initiation, at the end of the L1 stage (Fig 1.E). Indeed, as for a hs during embryogenesis, a late hs-mediated rescue in L1 is sufficient to fully suppress the defects. Rescuing LIN-15A activity after Y-to-PDA completion (in L3) on the other hand had no effect. Interestingly, a partial rescue could still be observed until the mid L2 stage. These could be indicative of a time-window in Td initiation: even though it starts just after the L1 stage, the process could still be rescued in a slightly larger time window by LIN-15A expression over the next hours.

### Two parallel initiation pathways

We previously reported that Y-to-PDA initiation was promoted by a plasticity cassette composed of a NODE-like complex - CEH-6/OCT, SOX-2/ SOX, SEM-4/SALL4, EGL-27/MTA1 - the EGL-5 Hox and the bHLH HLH-16 transcription factors (Kagias et al., 2012; Rashid et al., 2022; Daniele et al., 2025). As LIN-15A is equally required for the initiation step, we wondered whether it was acting in the same pathway as these factors.

First, we tested if *lin-15A* expression is under the control of the plasticity cassette. We found that endogenously tagged LIN-15A expression was unchanged in *sem-4(n1971)*, *egl-5(n945)*, *egl-27(ok1670)* or *hlh-16(fp12)* mutants compared to WT animals (Fig 2.A). These data suggest that *lin-15A* expression is not controlled by the previously described factors.

**Figure 2:**
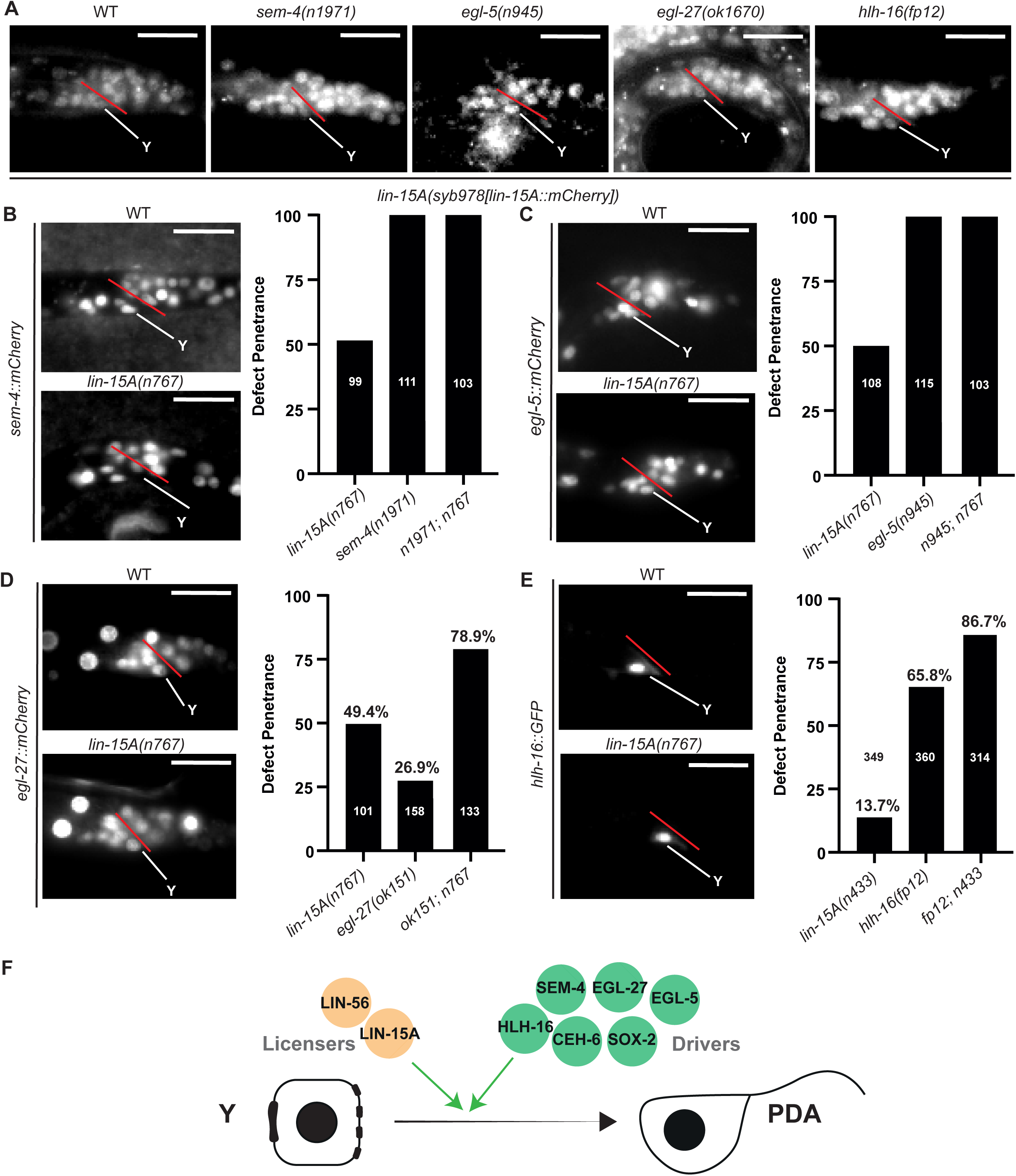
Two parallel pathways are required for Y-to-PDA initiation: Drivers and Licensers. In that entire figure: representative images of the rectal area during the L1 stage, prior to Y-to-PDA initiation. The rectal slit is indicated in red. Scalebars = 10 µm. **A** Expression of endogenous LIN-15A is not affected by the loss of any of the previously described factors *sem-4* (n=50), *egl-5* (n=47), *egl-27* (n=50) nor *hlh-16* (n=50) (for WT n=50). **B and C** LIN-15A acts independently of SEM-4 and EGL-5: SEM-4 and EGL-5 expression are not affected by the loss of *lin-15A* (for SEM-4 n WT=51 / n n767=48 – for EGL-5 n WT=52 / n n767=47) - Td defects induced by the loss of *sem-4* or *egl-5* are unchanged in double mutants with *lin-15A*. **D** LIN-15A acts independently of EGL-27: EGL-27 expression is not affected by the loss of *lin-15A* (n WT=48 / n n767=50) – *lin-15A(n767) (null)* induces defects in an additive manner with weak loss of function *egl-27(ok151).* LRT p-value (two-sided) between the *lin-15A* single and double mutants = 0.0034**. E**. LIN-15A acts likely independently of HLH-16: HLH-16 expression is not affected by the loss of *lin-15A* (n WT=50 / n n767=47) – *lin-15A(n433) (lf)* induces defects in an additive or slightly synergistic manner with strong loss of function *hlh-16(fp22).* LRT p-value (two-sided) between the *hlh-16* single and double mutants < 1 × 10⁻⁹. **F** The licensers (yellow) LIN-15A and LIN-56 act in parallel to the NODE-like plasticity cassette (Drivers - green).

Next, we tested whether LIN-15A could activate these factors. We found that the expressions of fluorescent reporters for SEM-4, EGL-5, EGL-27, HLH-16, CEH-6 and SOX-2 remain unchanged in *lin-15A(n767)* mutants compared to WT animals (Fig 2B-E and S2). We next used genetic interaction tests to assess the relationship between *lin-15A* and these factors. AS expected, the 100% penetrance of *sem-4(n1971)* and *egl-5(n945)* mutants remained unchanged in double mutants with *lin-15A(n767)* (Fig 2B and C). Double mutants between *lin-15A* and *egl-27* or *hlh-16* mutants that do not cause a full penetrance of Td initiation defects (Kagias et al., 2012; Zuryn et al., 2010; Daniele et al., 2025) showed additive effects, suggesting they act in different pathways (Fig 2D and E).

Together, our results show that LIN-15A and LIN-56 act in a distinct, novel pathway parallel to the plasticity cassette and therefore that multiple activities are required for successful Td initiation *in vivo* (Fig 2G): ‘Drivers’ (CEH-6/OCT, SOX-2/SOX, SEM-4/SALL4, EGL-27/MTA1 EGL-5/HOX and HLH-16/bHLH) factors that promote the dedifferentiation - whose loss result in total failure of initiation - and ‘Licensers’ (LIN-15A and LIN-56) that facilitate the process to happen - whose loss result in lower penetrance of Td initiation failure.

### LIN-15A acts as a (pro-)plasticity factor specifically in Y

We next tested if *lin-15A* is necessary in other plasticity events in *C. elegans*. We first tested whether it was required for the K-to-DVB transdifferentiation (Riva et al., 2022) (Fig 3Ai). K is another rectal cell that undergoes a transdifferentiation to produce a motor neuron named DVB which is critical for the proper defecation motor program (DMP) of the animal (Mahoney et al., 2008; Wang et al., 2013; Beg et al., 2008). In contrast to Y-to-PDA, K-to-DVB occurs through a cell division: K divides and its posterior epithelial daughter redifferentiates into DVB while the anterior daughter retains the rectal-epithelial identity. Nonetheless, the two Td exhibit similarities. Both Y and K are rectal cells and initiate their transdifferentiation at the end of the L1 stage. CEH-6/OCT, SOX-2/SOX2, SEM-4/SALL4 and EGL-5/HOX are also required for K-to-DVB, similarly to erase the rectal identity (Riva et al., 2022). Interestingly, LIN-15A is also expressed in K in L1 (Fig 3A ii) which further raised the question of its possible involvement in K-to-DVB. We found that neither LIN-15A nor LIN-56 were required for K-to-DVB (Fig 3B).

**Figure 3:**
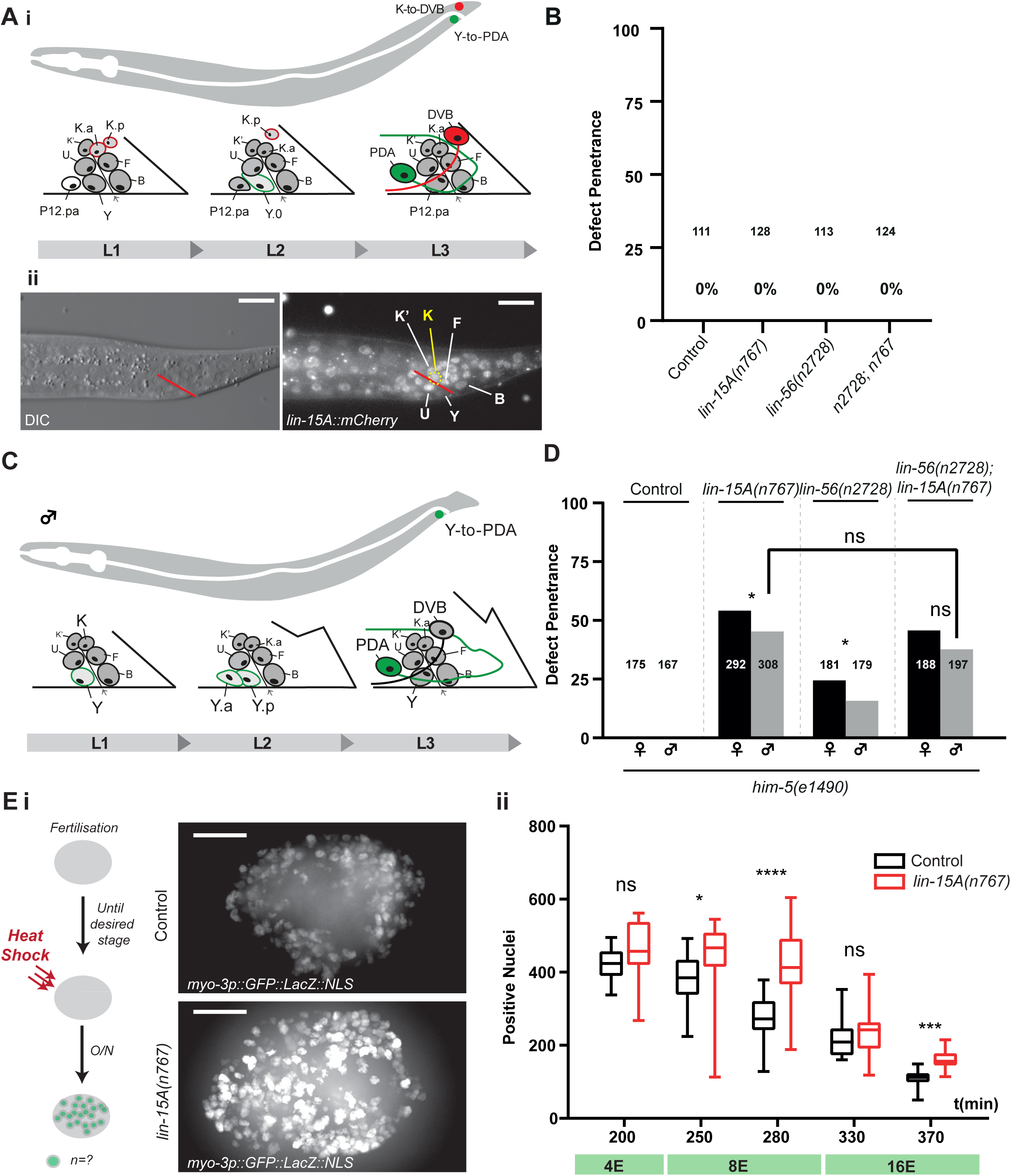
LIN-15A acts as a plasticity factor specifically in Y. **Ai** K-to-DVB transdifferentiation. At the end of the L1 stage, the K rectal cell divides. During the L2 stage the posterior daughter K.p redifferentiates into the DVB motor neuron. **ii** Images of the rectal area in L1 – prior to K-to-DVB Td. The rectal slit is indicated in red. LIN-15A is expressed in K (yellow) prior to its division. Scalebars = 5 µm. **B** *lin-15A* loss does not induce defects of K-to-DVB Td. Scorings performed in the absence or presence of DVB motor neuron. **C** Y-to-PDA transdifferentiation in males. A the late L1 – early L2 stage, the Y cell divides. At the late L2 / early L3 stage, the anterior daughter Y.a redifferentiates into the PDA motor neuron. **D** LIN-15A and LIN-56 act in cooperative manner during Y-to-PDA Td in males as they do in hermaphrodites. Scorings of male and hermaphrodite siblings from the same parental population and the same plates. Fisher exact tests with Bonferroni corrections for proportion comparisons. **E i** Number of converted blastomeres (*myo-3*-positive nuclei) at the five timepoints in control (black) and *lin-15A(n767)* animals (red) (the number of embryos scored at each time point is as follows : 200min wt=19 *n767*=10 - 250min wt=21 *n767*=20 - 280min wt=25 *n767*=17 - 330min wt=20 *n767*=19 - 250min wt=25 *n767=16*). In non-heat shocked WT animal, the number of body-wall muscle nuclei at birth is 95. Mann-Whitney tests with Bonferroni corrections for rank comparisons. **ii** Heat-shocked embryos displaying supernumerary *myo-3*+ blastomeres when HLH-1/MyoD is induced at 200, 280 or 370 minutes post-fertilization. To control that variations in the number of fluorescent nuclei was not caused by variations in the total number of nuclei, the total number of nuclei was monitored and remained constant in the different conditions. Examples of the z-stacks used for the 3D reconstruction and converted nuclei scorings are available at: https://drive.google.com/drive/u/1/folders/1TSElEi42NpMuBWCJ-KMi14RGqW4d0wFG.

As K-to-DVB involves a cell division, we wondered whether LIN-15A would be required in Y-to-PDA because of the lack of cell division. We therefore tested its involvment in Y-to-PDA in males, which occurs through a cell division (Riva C. and Jarriault S. *pers. comm*) (Fig 3C). Using the mutant *him-5(e1490)* which naturally produces around 35% of males within a population; we were able to score the presence or absence of PDA in males and hermaphrodites from the same mothers. We found that both LIN-15A and LIN-56 are required for Y-to-PDA in the males, and in the same non-additive and non-synergistic manner as in hermaphrodites (Fig 3D). Hence, the role of LIN-15A does not seem related to the absence of cell division, and may represent an event-specific feature of Y transdifferentiation.

We next examined whether it promoted early blastomere plasticity, especially as LIN-15A is widely expressed during *C. elegans* embryonic development (Fig S1.E) and is progressively lost as development advances. To test this hypothesis, we assessed whether loss of *lin-15A* impacts the Multipotency-to-Commitment Transition (MCT, reviewed in Rothman and Jarriault, 2019) which we found, using an improved protocol, to occur progressively from 200 to 370 minutes after fertilization (Lambert and Jarriault, *in prep*). We reasoned that if LIN-15A promoted embryonic cellular plasticity, the MCT should occur earlier in *lin-15A* mutants. We therefore performed a series of cell-fate challenge assays by forcing the expression of the muscle determinant HLH-1/MyoD, known to transform early embryos into a ball of muscle cells when expression is forced before the MCT (Coraggio et al., 2019; Fukushige and Krause, 2005; Yuzyuk et al., 2009). HLH-1/MyoD expression was overexpressed at 5 different time points during the timing of the progressive MCT (200/250/280/330/370 minutes post-fertilization) using a heat-shock promoter (Lambert and Jarriault, *in prep*). The strains used for the cell fate challenge assays carry an integrated array expressing an NLS::GFP under the control of the body-wall muscle-specific promoter *myo-3p*. Therefore, any cell differentiated into the body-wall muscle identity (both naturally and transdetermined) displays strong GFP signal (Fig 3E). Our results show that absence of *lin-15A* does not result in an earlier MCT, as would have been expected if it promoted early blastomere plasticity. Surprisingly, we even found that the loss of *lin-15A* increased the number of converted nuclei at 250 (early 8E stage), 280 (late 8E stage) and 370 (late 16E stage) minutes post-fertilization (Fig 3.E). In the control strain, the MCT clearly began between 200 and 250 minutes. In *lin-15A(n767)* mutants however, it is only first seen between 280 and 330 minutes. Overall, *lin-15A(n767)* blastomeres can be transdetermined for a longer period of time than their WT counterparts. Hence LIN-15A restricts the plasticity of the blastomeres rather than increasing it.

Together, these results highlight that LIN-15A does not increase cellular plasticity in general but acts as a pro-plasticity factor in specific cellular contexts like the rectal Y cell.

### LIN-15A antagonizes the SynMuvB chromatin modifiers which act as barriers for Y-to-PDA

We next examined the role of SynMuvBs genes, a class of genes known to act redundantly with *lin-15A* to repress *lin-3/EGF* during VPC specification (reviewed in Shin and Reiner, 2018). SynMuvBs code for members of chromatin complexes and modulators such as the DREAM, MEC or NuRD complexes (Table S1) and are major gene expression regulators (Fay and Yochem, 2007; Fay et al., 2002; Unhavaithaya et al., 2002; Wang et al., 2005; Petrella et al., 2011; Latorre et al., 2015; Wu et al., 2012; Käser-Pebernard et al., 2014; Goetsch et al., 2017; Ahringer and Gasser, 2018; Costello and Petrella, 2019; Rechtsteiner et al., 2019; Kipreos and van den Heuvel, 2019; Gal et al., 2021).

We reasoned that in the Y cellular context, the SynMuv genes could also act redundantly to LIN-15A. Under this hypothesis, we would expect single SynMuvB mutants to exhibit defects in Y-to-PDA Td, and/or double mutants *lin-15A + SynMuvB* to exhibit an increased defect penetrance (compared to the single *lin-15A* mutants). However, we found that no single SynMuvB mutant displayed defects in Y-to-PDA transdifferentiation. Moreover, the vast majority of the double mutants in combination with *lin-15A(n767)* actually displayed a defect suppression, from the typical 50% penetrance down to below 10% (Fig 4A-C). Thus, LIN-15A becomes dispensable when members of the SynMuvB factors are absent, suggesting that the latters act as a brake for Y-to-PDA Td, and are antagonized by LIN-15A. Accordingly, we found that defects of *lin-56(n2728)* were also suppressed in double mutants with *lin-35(n745)*, *lin-15B(n744)* and *lin-13(ok838)* (Fig S4A).

**Figure 4:**
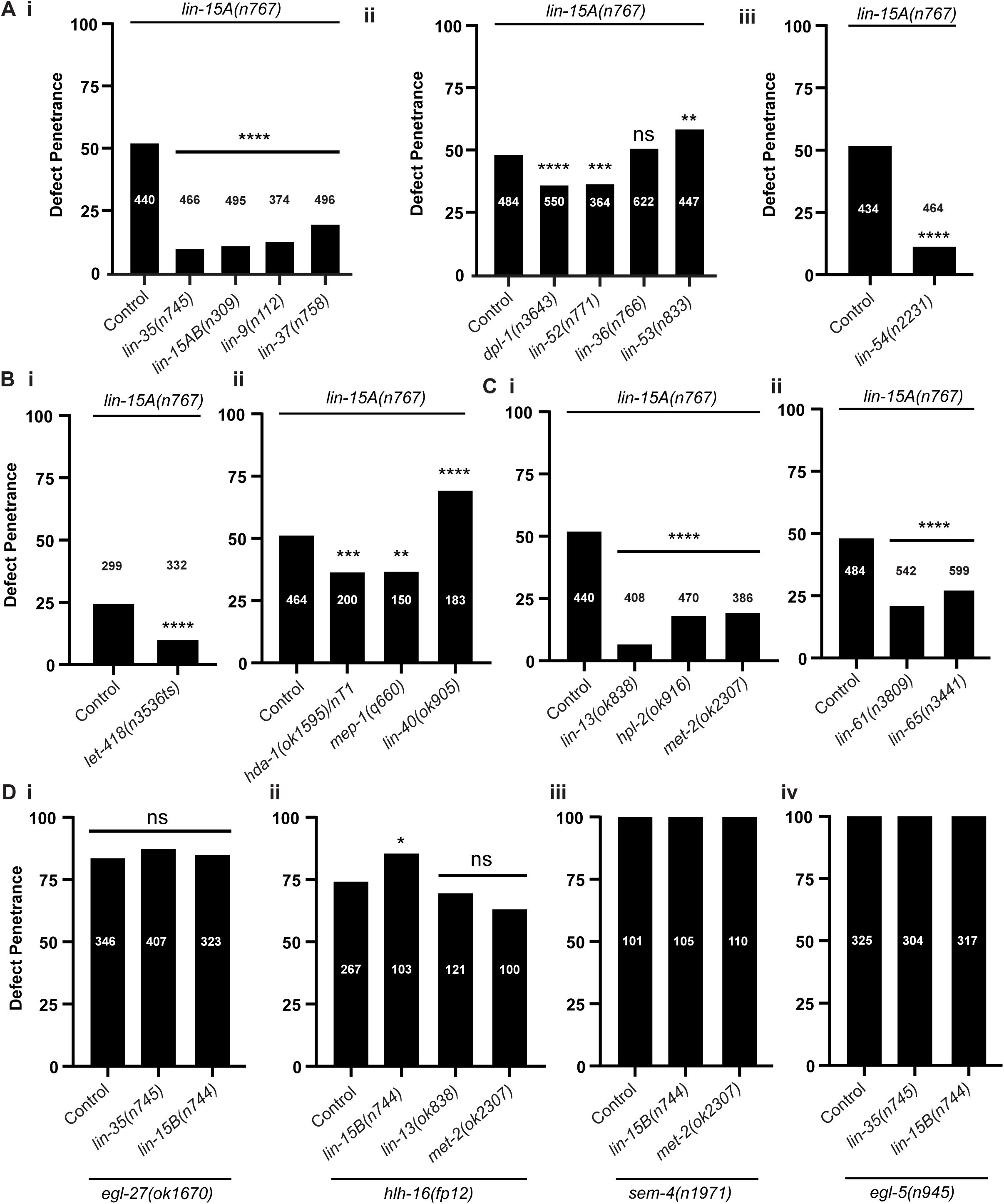
LIN-15A antagonizes the SynMuvB class of chromatin modifiers in Y-to-PDA. **A** Loss of members of the DREAM complex suppresses the defects of *lin-15A(n767)* mutants. X2 and Fisher exact tests with Bonferroni corrections for proportion comparisons. **B** Loss of the members of the NuRD and MEC complexes suppresses the defects of *lin-15A(n767)* mutants. *let-418(n3536)* is a temperature sensitive allele and was scored at 25°C. X2 and Fisher exact tests with Bonferroni corrections for proportion comparisons. **C** Loss of the SynMuvBs associated to the heterochromatin suppresses the defects of *lin-15A(n767)* mutants. X2 and Fisher exact tests with Bonferroni corrections for proportion comparisons. **D** Losses of SynMuvB chromatin modifiers do not suppress Td defects induced by the loss of the Driver factors EGL-27(**i**), HLH16 (**ii**), SEM-4 (**iii**) nor EGL-5 (**iv**). X2 and Fisher exact tests with Bonferroni corrections for proportion comparisons.

As the function of the SynMuvBs and LIN-15A (as a SynMuvA) in VPC specification is to repress *lin-3/EGF*, and as the Y cell is rectal-epithelial, we also tested whether *lin-3/EGF* is involved in Td initiation. We assessed the effects of the loss-of-function alleles of *lin-3/EGF mt378* and *e1417*, and of the gain-of-function allele *n4441* (Saffer et al., 2011), alone and in double mutants with *lin-15A(n767)*. In *n4441* a mutation in the promoter of *lin3/EGF* prevents its repression by the SynMuvAs including LIN-15A. We found that none of these *lin-3/EGF* alleles induced defects on their own, nor did they alter defect penetrance in *lin-15A(n767)*. Hence, the functions of LIN-15A and of the SynMuvBs in Y-to-PDA are independent of *lin-3/EGF*.

### The two NuRDs complexes of *C. elegans* have opposite effects on Y plasticity

Out of all the SynMuvB genes tested, two stand out since their losses did not suppress the defects of *lin-15A(n767)* mutants (Fig 4A): loss of *lin-36* had no effect while loss *lin-53* further significantly enhanced Td defects from 50% up to 65%. The absence of LIN-36 involvement in Y-to-PDA postmitotic Td (Jarriault et al., 2008) is consistent with its repression of genes associated with cell cycle progression (Gal et al., 2021).

However, loss of *lin-53* unambiguously increases the defects of *lin-15A* mutants. LIN-53 is part of several complexes including the NuRD complex, as are *hda-1* and *let-418* which loss result in a decrease of *lin-15A(n767)* defects (Fig 4B). We found that the loss of another core component of the NuRD complex, *lin-40*, which is not a SynMuvB, also increased the defects of *lin-15A* mutants. Two distinct NuRD complexes co-exist in *C. elegans* (Käser-Pebernard et al., 2016; Zelewsky et al., 2000), which differ by their Mi2 homolog, either LET-418 (a SynMuvB) or CHD-3 (Fig S5A). We hypothesized that the opposite impact of loss of NuRD components on *lin-15A(0)* defects could be explained by different activities of the 2 NuRDs. Therefore, we specifically tested whether the two NuRD complexes could have opposite effects on cellular plasticity. Under this hypothesis, we predicted that if NuRD*LET*−418 restricts Y plasticity – as *let-418* loss suppresses *lin-15A(n767)* Td defects, then NuRD*CHD*−3 would further enhance it. As hypothesized, we found that the loss of *chd-3* increased Td defects in *lin-15A(n767)* mutants, similarly to the loss of *lin-40* or *lin-53*, while having no effect in WT backgrounds (Fig S5B). Thus, our data suggest the existence of two NuRD complexes with distinct roles in Y-to-PDA Td. These data are in accordance with previous reports of their distinct and sometimes opposite activities in other contexts (Passannante 2012; Käser-Pebernard 2016).

Interestingly, CHD-3 loss also enhanced the defects observed in *egl-27(ok151)* mutants (Fig S5C). Since LIN-15A and EGL-27 act in two parallel pathways, these results suggest that the action of the NuRD*CHD*−3 complex is not specific to the Licensers/SynMuvBs nor the Drivers but act in a similar manner on both pathways (Fig S5D). Thus, Y-to-PDA involves a complex dance of various and close chromatin complexes that either facilitate it or block it.

### The SynMuvBs specifically interact with the Licensers

Next, we examined if the SynMuvBs were only antagonized by the Licencers LIN-15A and LIN-56, or if they also interacted with the Drivers (Fig 4D). We tested if loss of the SynMuvBs *lin-35*, *lin-15B*, *lin-13* or *met-2* suppressed Td defects in Drivers mutant *egl-27(ok1670)*, *hlh-16(fp12)*, *sem-4(n1971)* and *egl-5(n945)*. No significant suppression was found in any combination (Fig 4D) which confirmed that the SynMuvBs and LIN-15A act antagonistically in the same pathway or on the same targets, and so that the function of LIN-15A lies in its interaction with the various SynMuvBs. Furthermore, this strengthens our model of two independent pathways (Fig 5).

**Figure 5:**
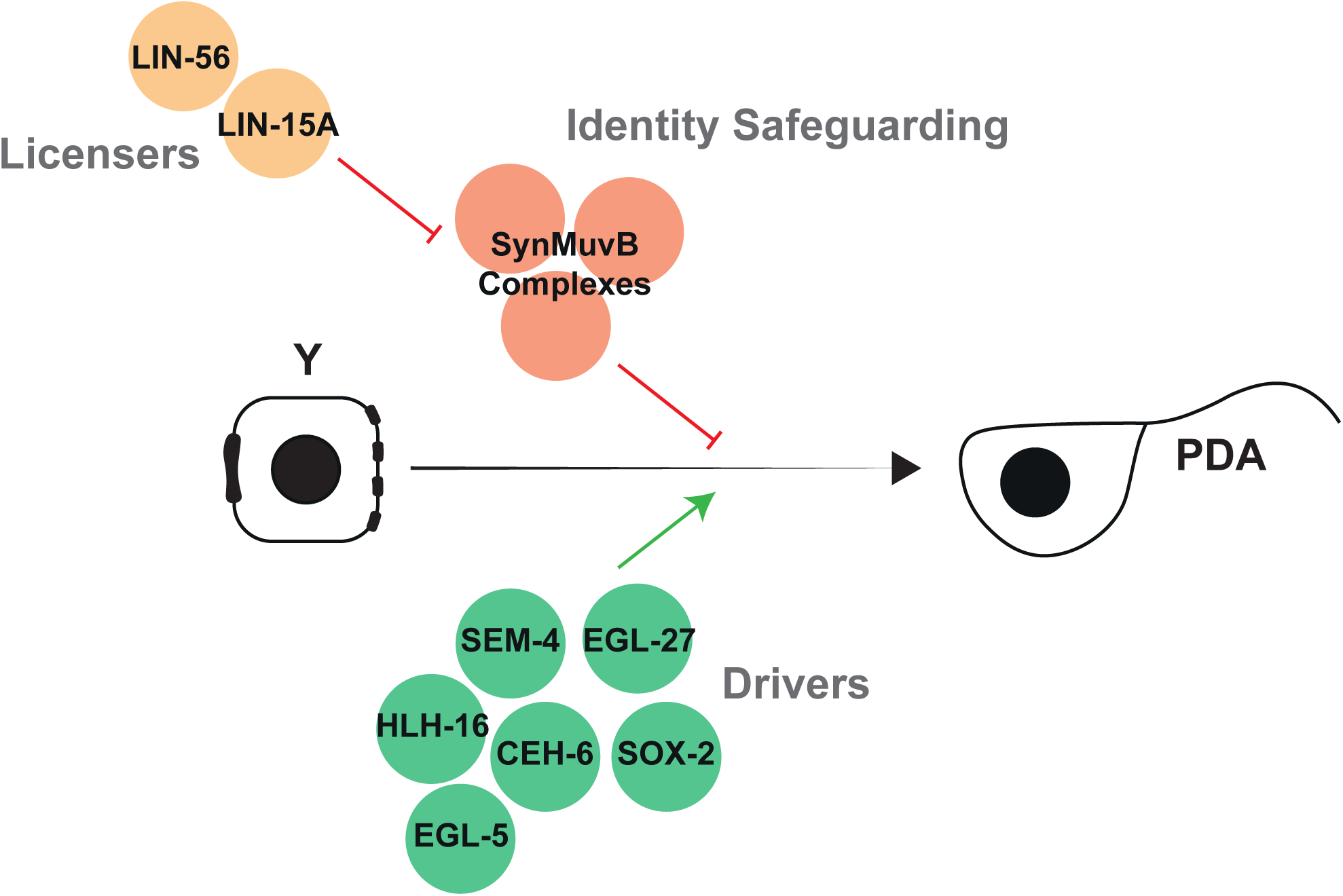
The Licensers (yellow) LIN-15A and LIN-56 antagonize mechanisms that safeguard the initial rectal-epithelial identity (red). In parallel, Driver factors (green) trigger Td initiation.

### The SynMuvBs mainly act in the rectal cells for their role of barriers for Y-to-PDA transdifferentiation

To further address the role of the SynMuvBs in Y-to-PDA and their relationship with LIN-15A, we chose to focus on LIN-35, the sole *C. elegans* ortholog of pRb and core component of the SynMuvB-containing DREAM complex. We reasoned that since most display a similar genetic interaction with *lin-15A*, it is likely that they act coordinately against Y-to-PDA initiation. This paradigm is supported by previous work reporting that the binding regions of numerous SynMuvBs extensively overlap (McMurchy et al., 2017) and that the DREAM complex represses its targets by depositing the repressive mark H3K9*me*2 thanks to the SynMuvB MET-2 (Gal et al., 2021).

We first sought to identify in which tissue the lin-35 was required for its role in Y-to-PDA initiation. Hence, we performed tissue-specific rescue experiments of LIN-35/pRb in *lin-35(n745); lin-15A(n767)* animals (Fig S4B). We found that rescuing the expression of *lin-35* ubiquitously (under *eft-3p* promoter) or in the rectal area (under *egl-5p* promoter) consistently restored Td defects (Fig S4B). In contrast, intestinal (under *ugt-22p* or *elt-2p* promoters) and neuronal (under *rab-3p* promoter) expression led to much more limited and inconsistent rescues (Fig S4B). Given these results, we concluded that LIN-35/pRb mainly acts in the rectal cells during Y-to-PDA. However, it may have another role in non-rectal tissues that contribute – to a much lesser degree – to its function as a brake for Td.

### LIN-15A does not affect SynMuvB expression in Y

We next sought to understand how LIN-15A antagonizes LIN-35/pRb and MET-2 in Y-to-PDA transdifferentiation. The first possibility is that LIN-15A represses their expression, or their nuclear localization. We compared the expression and the localization of endogenously tagged LIN-35/pRb and MET-2 in WT animals and in *lin-15A(n767)* mutants. Both proteins are strictly nuclear as previously described by Gal et al. (2021); Delaney et al. (2019), and we never observed any change in their localization. Because we showed that LIN-35/pRb was crucial in the rectal cells for its role in Td, we measured the fluorescence intensity within the nucleus of the Y cell. We found that the expression of LIN-35/pRb was not altered by the loss of *lin-15A* (Fig S4C) and no obvious change was detected in any of the other rectal cells either. We even found that the MET-2 signal was slightly lower in *lin-15A(n767)* mutants compared to the WT (Fig S4D), while the opposite would be expected if LIN-15A repressed *met-2* expression. Hence, the antagonization of the SynMuvBs by LIN-15A is not likely mediated by a negative control of their expression but rather by impacting their activity.

### LIN-15A modulates the binding of the DREAM complex

To better understand the functional antagonism between the DREAM complex and LIN-15A, that both bind chromatin (Latorre et al. 2015; Gal et al. 2021) (Fig S6), we sought to compare their targets and dependencies through Chromatin Immuno-Precipitation (ChIP)-seq experiments, using antibodies against the endogenous LIN-35, LIN-15B and LIN-15A on WT animals, *lin-15A(fp31)* (CRISPR knock-out of entire *lin-15A* locus), *lin-15B(n744)* (null) and *lin-35(n745)* (null) mutants. The DREAM complex represses cycle genes in combination with THAP domain protein LIN-36, and represses germline genes in the soma in combination with THAP domain protein LIN-15B and H3K9*me*2 promoter marking (Gal et al., 2021). Given that LIN-15A possesses a THAP domain required for its role in Y-to-PDA (Fig 1.D) and binds onto naked and nucleosome-bound DNA (Figure S6), we reasoned that LIN-15A also acts on chromatin, and might modulates DREAM activity. We performed ChIP-seq in early stage, unstarved, L1 larvae as it was not technically feasible to purify sufficient Y cells for analysis.

As previously reported for starved L1s (Gal 2021), LIN-35 and LIN-15B had similar binding patterns and the vast majority of those peaks are located in promoter regions (SI Fig 6C). We found 5382 and 6540 peaks bound by LIN-35 and LIN-15B respectively, with 65,3% shared genes (n=2911) in WT animals. These targets are consistent with previous data obtained from L1 larvae synchronized through starvation (95,5% and 90% overlap for LIN-35 and LIN-15B respectively, Table S2), and correspond mostly to metabolic processes (biogenesis, biosynthesis), developmental and cell cycle-related (DNA replication and mitotic) BP GOs (Table S2). LIN-15A was found to bind 5295 regions, the vast majority of which are also located in promoter regions (SI Fig 6C). LIN-15A bound genes are enriched with similar BP and CC GO terms as LIN-35 and LIN-15B bound genes and KEGG hits related to metabolic and basal developmental pathways (TOR, IIS, TGF-*β* or Wnt, Table S2). 73.9% of LIN-15A-bound genes (n=3340) are in common with LIN-15B and 66.3% (n=2996) with LIN-35-bound genes (SI Fig 6D), supporting changes in the chromatin state of specific loci as the outcome of LIN-15A-DREAM antagonist interaction. Furthermore, most of the genes common between LIN-15A and LIN-15B or LIN-35 are actually bound by all 3 factors (61.3%, 2770 genes; Figure S6D).

**Figure 6:**
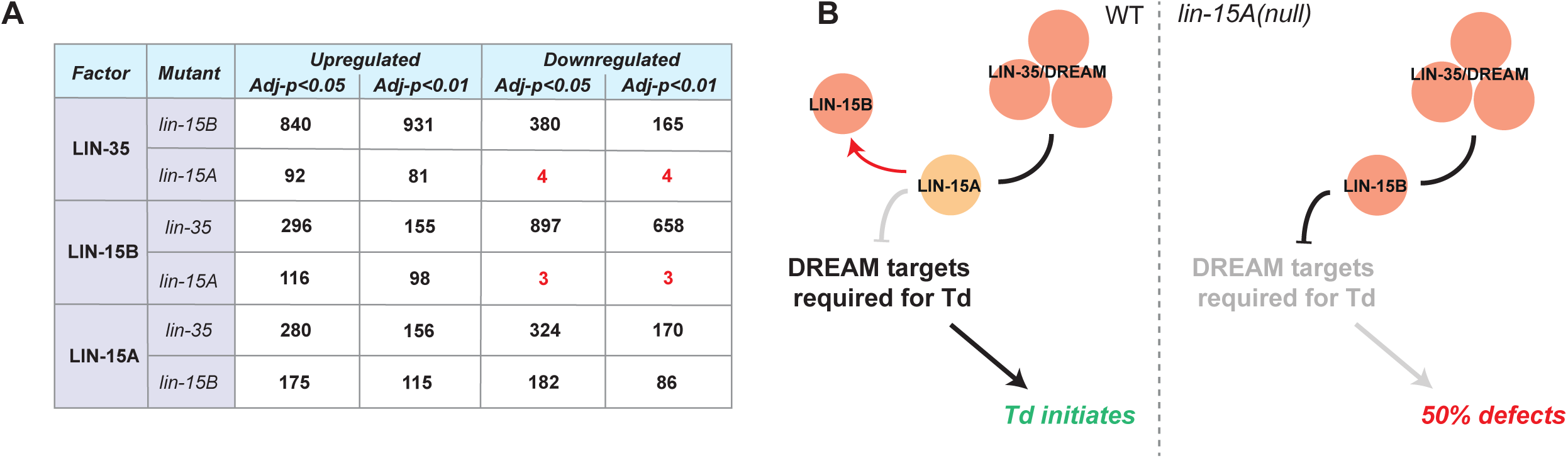
LIN-15A binds chromatin and can modulate DREAM binding onto a subset of its targets. **A** Number of peaks differentially bound by LIN-35, LIN-15B and LIN-15A in the different mutants. Loss of *lin-15A* increases the binding of LIN-35 and LIN-15B much more than it decreases it suggesting that LIN-15A only prevents DREAM binding, on a subset of its target genes. In contrast, losses of *lin-15B* or *lin-35* in have mixed effects on LIN-35 or LIN-15B binding. Similarly, the losses of *lin-35* or *lin-15B* have mixed effects on LIN-15A binding. **B** Proposed molecular mechanism of LIN-15A, LIN-35 and LIN-15B interaction in Y-to-PDA. LIN-15A may shield certain DREAM targets relevant for Td from its gene repression activity. In *lin-15A* mutants, those targets are not protected from DREAM repression, and get turned off, which impairs Td initiation.

To ask whether the binding of LIN-15A, LIN-35 or LIN-15B are impacted by each other, we analyzed whether peaks change in mutants. We observed that hundreds of LIN-35 or LIN-15B peaks had reduced or increased signal when the other factor was absent. Similarly increased and decreased LIN-15A peaks were observed in absence of LIN-13 or LIN-15B. In contrast, loss of LIN-15A led to increased signal for ∼100 LIN-35 or LIN-15B peaks (Figure 6A and Table S2). Absence of *lin-15A* specifically led to an increase in occupancy as very little binding loss is observed for these factors in *lin-15A* mutant (Figure 6A and Table S2). We found no significative GO enrichment for the small sets of LIN-35 (n=85) and LIN-15B (n=105) bound genes due to their limited numbers. However, the Groups tool from ShinyGo shows that many of the genes where LIN-35 or LIN-15B bindings are increased in *lin-15A(fp31)* mutants fall into CC GO groups related to membrane and vesicle trafficking (Table S2). Such biological functions are reminiscent of the CC GOs found for the shared genes bound by all three factors in WT animals, and could suggest that membrane processes could play a key role in initiating transdifferentiation. Altogether, these data suggest that LIN-15A activity prevents DREAM binding to a subset of genes.

## Discussion

### Drivers and Licensers

Specific differentiated cells maintain the ability to change their identity even at later stages of development, in a process called natural transdifferentiation (Lambert et al., 2021; Okada, 1991; Eguchi, 1993). In *C. elegans*, the rectal Y cell, that transdifferentiates into the PDA motoneuron, is a well-characterized example (Jarriault et al., 2008; Richard et al., 2011; Lambert et al., 2021). We previously found that several factors, together designated as ‘plasticity cassette’, are required for the initiation step (Kagias et al., 2012; Daniele et al., 2025).

In this study, we have found that a second pathway centered on the SynMuvA LIN-15A is also required for Y-to-PDA. LIN-15A, a nuclear factor able to bind chromatin, acts in a cooperative manner - likely forming a dimer - with LIN-56, to enable Td initiation. Importantly, LIN-15A and LIN-56 act in parallel to the previously described factors. We further found that the Td defects in *lin-15A* mutants, but not in plasticity cassette mutants, could be suppressed by the loss of most SynMuvBs (Fig 4 and S4). Altogether, our genetic data argue for LIN-15A, LIN-56 and SynMuvBs to act antagonistically to each other and in parallel to plasticity cassette factors. Hence, we propose a two-pathway model (Fig 5): on the one hand, conserved plasticity factors (NODE-like complex) are absolutely required to dedifferentiate the rectal Y cell, acting as ‘Drivers’. Their losses result in 100% (or close to 100%) defects in initiation. On the other hand, LIN-15A and LIN-56 are required to facilitate the initiation of Y-to-PDA (their loss does not induce full Td defect penetrance), through antagonization of SynMuvB activity, therefore acting as ‘Licensers’. Several questions arise from these results. Why does SynMuBs activity represent a barrier for Y-to-PDA transdifferentiation? Are they general brakes to reprogramming? How does LIN-15A antagonize then in Y-to-PDA?

### The SynMuvBs as cell identity maintenance mechanism

The SynMuvBs, which likely act in the Y cell, do not appear to impact on the Y cell identity itself: Y is formed in all examined SynMuvB mutants and the expression of rectal markers is unaffected by the loss of *lin-15A* (Fig S1D). Hence, their activity likely only provides robust maintenance of the differentiated Y ID once established.

SynMuvBs have been described as chromatin factors, the majority of which are part of different multimeric complexes, and act to enforce an inactive chromatin state. We suggest that these chromatin factors are repressing, in the Y cell, the expression of genes associated with other fates, thus re-enforcing Y rectal identity by default. De-repression of one or more alternative transcriptomes would allow progression towards an alternative fate, here neuronal. Such role would be consistent with their role in other cellular contexts (Ahringer and Gasser, 2018; Goetsch et al., 2017; Latorre et al., 2015; Unhavaithaya et al., 2002; Gal et al., 2021; Fay et al., 2002; Wang et al., 2005; Wu et al., 2012; Käser-Pebernard et al., 2014; Rechtsteiner et al., 2019; Kipreos and van den Heuvel, 2019). For instance, SynMuvBs antagonize germline fate in the intestine (Petrella et al., 2011). In brief, the role of the Licensers would be to provide a favorable chromatin context for cellular processes that favor/install a plastic state, possibly through the modulation of membrane processes as suggested by our ChIP-seq analyses (Fig 6). On the other hand, the Drivers role would be to activate Td genes or inactivate rectal genes. Another non-exclusive hypothesis is that SynMuvBs could repress the genes that make the Y cell plastic and able to initiate the Td process.

### The SynMuvBs as barriers for cellular reprogramming

Our result on this SynMuvB brake on Y-to-PDA natural Td parallels the need to remove chromatin modifiers to enable induced reprogramming both in *C. elegans* (Tursun et al., 2011; Patel et al., 2012; Hajduskova et al., 2019; Kolundzic et al., 2018) and in other systems (reviewed in Brumbaugh et al., 2019, Rothman and Jarriault, 2019). In *C. elegans* notably, knockdown of SynMuvB LIN-53/RbAp48 by RNAi is required to successfully induce transdifferentiation of germ cells into neurons upon CHE-1 forced expression (Tursun et al., 2011). Thus, similar barriers impede Td in both natural and induced settings, although some context-specificity is observed: LIN-53/RbAp48 promotes, alongside LIN-15A, the natural transdifferentiation of the Y cell while it blocks induced reprogramming of germ cells. Thus, the nature of the interactions between these chromatin modifiers and the precise cocktail safeguarding a given differentiated identity is likely specific to the considered cell.

### Context specificity of the Licensers

Our data suggest that Licensers activity is context-dependent. Indeed, we found that LIN-15A is required for Y-to-PDA transdifferentiation both in males and hermaphrodites, independently of cell division (Fig 3D). However, while also expressed in K, LIN-15A is not required for K-to-DVB, despite occurring in the same organ, at the same time, and requiring members of the NODE-like complex (Riva et al., 2022). Furthermore, in some other contexts such as embryonic development LIN-15A can even restrict cellular plasticity (Fig 3F). Hence its role to promote plasticity appears specific to certain cellular identities, and this is paralleled with the changing nature of its interactions with SynMuvBs. Likewise, LIN-53 has been shown to block induced reprogramming of the germ cells (Tursun et al., 2011) while it promotes natural transdifferentiation of the Y cell alongside Lin-15A. One possibility is that being plastic entails very different requirements in different cells, dictated by the nature and the sturdiness of the brakes to remove, or the type of transcriptional program to switch on and/or the extend of the necessary alterations to different programs. Alternatively, while LIN-15A acts as a Licenser in the Y cell it is possible that other factors play a similar role in other contexts. Work from the Hobert and Tursun labs previously highlighted the diversity of factors that block induced reprogramming (Rahe and Hobert, 2019; Tursun et al., 2011; Patel et al., 2012; Kolundzic et al., 2018; Hajduskova et al., 2019). While the control of chromatin remodeling and TF activity are key recurring themes, the precise identity of the players involved clearly varies depending on the cellular context. The diversity of chromatin modifiers available to *C. elegans* allows for modular flexibility: some complexes and interactions may be required for instance to safeguard certain identities, while other combinations would have different functional outcomes. Our findings that the two *C. elegans* NuRD complexes exhibit antagonistic activities with regards to Y-to-PDA illustrate this fluidity not only in complex composition but also in functional outcomes. LIN-15A is another prime example, acting in a redundant manner with the SynMuvBs – and in combination with other SynMuvAs - to repress *lin-3/EGF* for correct VPC specification, but that acts as a pro-plasticity factor in Y-to-PDA by antagonizing the SynMuvBs, and without other SynMuvAs (apart from LIN-56).

### The SynMuvBs may act as a single machinery in Y-to-PDA

We found that LIN-15A antagonizes all of the SynMuvBs, except LIN-53 because of its role in the two alternative NuRD complexes (Fig 4A, 4B and S5). This suggests that the SynMuvBs act as a single multimodal machinery in Y-to-PDA. Supporting this idea, work from the Ahringer lab has highlighted that the binding regions of LET-418/Mi2, HPL-2, LIN-61, LIN-13 and MET-2 - all involved in antagonizing Y-to-PDA - extensively overlap (McMurchy et al., 2017), and that LIN-35 and LIN-15B repress their targets via H3K9me2 deposition by MET-2 (Gal et al., 2021). Such interactions between SynMuvB complexes have also been reported in mammalian cells, where Sin3B, a protein associated with HDAC (orthologue of NuRD component HDA-1) interacts with the DREAM and is required for repression of its targets to promote cell quiescence (Bainor et al., 2018). Hence, LIN-15A may antagonize a single machinery involved in gene repression. Supporting this view of regulation at the chromatin level, LIN-15A THAP domain – for DNA binding – is required for its role in Y-to-PDA (Fig 1D and S6).

### The antagonism between LIN-15A and the SynMuvBs may take place at the chromatin level

Since the SynMuvBs may act as a single machinery in Y-to-PDA, we focused on the possible interaction between LIN-15A and the DREAM complex. Previous studies highlighted that the latter acts through THAP domain proteins, notably LIN-15B (Latorre et al., 2015; Gal et al., 2021). Since LIN-15A is in an operon with LIN-15B, also possesses a THAP domain, required for its role in Td, and can bind DNA (Fig 1.D and S6), we reasoned that it could also interact with the DREAM, hence explaining at the molecular the genetic antagonization. We found that the binding regions of LIN-35/DREAM, LIN-15B and LIN-15A to vastly overlap and to favor promoters (Fig 6A and 6B). These results are consistent with previous records of LIN-35 and LIN-15B binding (Gal et al., 2021). As LIN-35/DREAM and LIN-15B repress gene expression via MET-2 H3K9*me*2 promoter marking, these results further suggest a modulation of gene expression through chromatin remodeling by LIN-15A.

In accordance with this hypothesis, our ChIP data from mutant backgrounds suggest that LIN-15A prevents LIN-35/DREAM and LIN-15B binding to a subset of their target, which may alleviate their repression. Therefore, by limiting binding to these genes, LIN-15A activity could prevent DREAM-mediated H3K9*me*2 promoter marking, thus alleviating expression repression on Td-relevant targets. GO terms associated with a large share of these targets are notably associated with membrane processes (Table S2). While our ChIP data were obtained at the whole organism level and may mask event-specific binding, it is interesting to note that the first materialization of Y transdifferentation is its retraction from the rectal tube which, at the molecular level, is correlated with a *sensu-strictu* dedifferentiation (Jarriault et al., 2008; Richard et al., 2011). Overall, this process resembles an epithelial-to-mesenchymal transition (EMT), a process that involves important membrane remodeling (Jarriault et al., 2008; Ferreira et al., 2024; Khan and Steeg, 2021; Guo et al., 2024; Belenyesi et al., 2025). Interestingly, induced EMT of mammary epithelium in mice has been shown to results in expression of stem cell properties (Mani et al., 2008). Thus, retraction of the Y cell from the rectum, which is reminiscent of an EMT, could be a plasticity triggering event rather than a consequence. However, the EMT process has not been characterized *per se* in *C. elegans* and mutants of *C. elegans* homologues of EMT effectors (e.g. snail1, 2, twist or eomes) do not exhibit Y-to-PDA defects (S. Zuryn, SJ, personal communication). It is possible that delamination of the Y cell from the rectal tube relies on other, uncharacterized, mechanisms. Overall, an intriguing possibility is that the action of LIN-15A would be to increase the delaminating capacities of Y by alleviating SynMuvB repression of membrane processes genes, and that this event would increase the ability of the Y cell to swap identity.

### Antagonization of pRb in the context of reprogramming

LIN-35 is the sole pocket protein encoded in the *C. elegans* genome. Its mammalian counterpart, the retinoblastoma protein (pRb), is a major tumor suppressor, whose loss plays a critical role in the development and growth of multiple cancers. Most of the functions of pocket proteins - notably cell cycle control and differentiation - are conserved throughout eukaryotes, including vertebrates, nematodes, and even plants (Zluhan-Martinez et al., 2020; Desvoyes and Gutierrez, 2020). Importantly, one of the major anti-oncogenic roles of mammalian pRb is to inhibit cancer cell EMT (Egger et al., 2016)). Thus, it might be that *C. elegans* LIN-35 retained a similar function, preventing the acquisition of migratory phenotypes for differentiated cells. pRb has also been shown to repress the reprogramming of murine fibroblast into iPSCs (Kareta et al., 2015). Y-to-PDA transdifferentiation appears to represent another, conserved role of pocket proteins in regulating reprogramming. Kareta and colleagues equally showed that this function of pRb was independent on its role in regulating the cell cycle, data reflecting our own report that LIN-36 - the DREAM adaptor mediating LIN-35/pRb repression on cell cycle-associated genes (Gal et al., 2021), does not play any role in Y Td. Altogether, these findings suggest that pocket proteins may have a conserved ability to restrict cellular plasticity, independently of their control of the cell cycle.

### Limitations of this study

As no Y-specific driver is known to this day, we cannot exclude that LIN-15A and/or LIN-35 act in the other cells where the rectal-specific *egl-5p* promoter also drives expression. However, given the roles in gene expression regulation, nuclear localization and DNA binding of those factors, this alternative hypothesis is much less probable than cell-autonomous roles in Y.

The ChIP-seq performed in this study could only be carried out at the whole animal level. Hence, additional relevant targets involved in Y plasticity may have been missed. This limitation comes from technical reasons. The epithelial-rectal Y is wrapped in the resistant L1 cuticle and relatively small - approximately 5 µm – making the purification of large amounts (*>*3000 individual cells) difficult.

For ChIP-seq experiments, we focused on the interaction between LIN-15A and the DREAM complex based on our genetic data (Fig 4), and on reports that the DREAM acts notably through LIN-15B, on a operon with LIN-15A and sharing similar THAP domains (Gal et al., 2021). We reasoned that SynMuvBs act as a single machinery in Y-to-PDA (see above), but we cannot exclude that LIN-15A and other SynMuvBs interact independently of LIN-35.

## Materials and Methods

### *C. elegans* culture, Y-to-PDA scorings and general experimental procedures

*Caenorhabditis elegans* strains were maintained according to standard protocols (Brenner, 1974) - 20°C, OP50 on Nematode Growth Media (NGM) agar plates. For all experiments, a parental population was amplified. Broad synchronization was performed by hypochlorite treatment (bleach), embryos were next placed on fresh plates (max 3 days old), seeded with fresh bacterial lawns (1 day old O/N liquid culture) dried after seeding at 37°C for a few hours. OP50 loans were grown from single colony to saturation in Luria-Bertani Broth (LB) at 37°C, shaking at 170rpm. This procedure was found critical to have consistent penetrance of the *lin-15A* mutants. Presence of absence of the PDA motor neuron (scoring) were performed at the late L3, L4 or young adult stage. Defects were defined by the absence of PDA (no cell body, no axon, no marker expressed). PDA presence was assessed with the integrated array *syIs63 (cog-1p::GFP)* or *wyIs75 (exp-1p::GFP)*. Defect penetrance does not depend on the marker used for mutants involved in the initiation step. K-to-DVB was scored in a similar fashion, using the integrated array *wyIs75 (unc-47p::RFP)*. Experiments were conducted in triplicates at minima. Except stated otherwise, all experiments were conducted at strictly 20°C, with OP50 as a food source on NGM plates. Scoring of *let-418(n3536)* was notably conducted differently. *n3536* is a temperature sensitive allele of *let-418* behaving as a null at 25°C (C. Wicky *pers. comm*). At 25°C however, *n3536* leads to a developmental arrest. In order to use *n3536* but leave time for Y-to-PDA to proceed, without being impact by the developmental arrest, synchronized newly-hatched larvae were placed at 20°C and transferred at 25°C 6 hours later. This procedure allows *let-418(n3536)* to develop reaching the L3 and L4 stages. The efficiency of this procedure was confirmed by the induction of the Multivulva phenotype in *lin-15A(n767); let-418(n3536ts)* mutants, the irreversible developmental arrest and subsequently the reduced penetrance of Y-to-PDA defects. Of note, the Td defect penetrance of *lin-15A(n767)* mutants is decreased at 25°C compared 20°C from 50% down to 25%.

For epifluorescence microscopy, animals were immobilized on % agarose pads using Tricaine 0.4% and Tetramisole 0.04% in UP water. Leica DM6 B microscope was used, and images were captured with LAS X software through Hamamatsu Digital Camera C11440. Standard image analyses were performed with FIJI.

*C. elegans* transgenic strains were engineered by DNA microinjection into the gonad cytoplasm of young adults (Mello et al., 1991). The mix of injected DNA was always composed by the plasmid of interest together with a co-injection marker and pBSK for a final concentration of DNA of 200-250 ng/µl in water or injection buffer (2% polyethylene glycol MW 6000-8000, 20 mM potassium phosphate pH 7.5, 3 mM potassium citrate pH 7.5). Transgenics F1 were cloned and the transmission of the plasmid array was evaluated in the F2 generation. Lines with a good transmission rate were usually kept. All the other strains used for this study have been obtained by crossing existing strains from our lab, other labs (obtained through CGC or directly) or from SunyBiotech for KI reporter strains which have been designed in the lab. The presence of mutations was always confirmed prior to the experiments by PCR genotyping in case of deletions and by PCR plus restriction digestion genotyping in case of point mutations (Morin 2020). Lysates of entire worms were produced for genotyping. Worms were lysed in a solution containing ultrapure water, PCR buffer 1X and proteinase K 0.5 mM, for 1 h at 65°C and then at 95°C for 15 min.

### Statistical analyses

Statistical analyses were performed with PRISM GraphPad 10. For all experiments, the statistical tests used are indicated in the corresponding figure legend. ns: p*>*.0.05, *: p*<*0.05, **: p*<*0.01, ***: p*<*0.001, ****: p*<*0.0001.

### Isolation and cloning of ***fp22***

*fp22* was isolated from an EMS screen as a “no PDA” mutant by absence of the expected staining by *syIs63* transgenic marker. The mutant was backcrossed and EMS variant deep sequence mapping was performed as described in (Zuryn et al., 2010).

### RNA interference

For *lin-8*, *lin-38*, *smo-1* and *uba-2*, a gene fragment from a cDNA library made from RNA extracted from N2 worms was PCR amplified with primers containing flanking T7 promoter sequences. In vitro transcription using the PCR products as templates and T7 RNA polymerase was performed, and single stranded RNA was purified using Qiagen RNeasy Mini Kit (74104) and allowed to anneal as dsRNA by gradually lowering the temperature of the sample. dsRNA was injected into *rrf-3(pk1426); syIs63* or *rrf-3(pk1426); syIs63; lin-15A(n767)* adults and F1 progeny derived from these adults were scored for Y-to-PDA defects.

### Tissue specific, truncated domains and heat-shock inducible rescues of LIN-15A and LIN-35

Coding sequence were cloned from a complementary DNA (cDNA) library prepared from WT (N2) RNA that had been extracted using RNA-Solv. Promoter sequences were cloned from WT (N2) DNA. All final constructs were sequenced verified before use.

### *lin-15A* rescue constructs

The *lin-15p::mcherry::lin-15A cDNA* transgenic plasmid pSJ805 contains the *lin-15* promoter derived from N2 DNA via PCR using the primers F: 5’- GGTCGACCGGCCGACCGTTTTCTTTCG-3’ and R: 5’- GGGGATCCAATTACCTGAAAATATAAAAAAAATAATAAATTA-TTCGGTCATCATGC-3’ and cloned in between the Acc1 and MscI sites of pPD49.78 (Addgene plasmid 1447; http://n2t.net/-addgene:1447; RRID:Addgene 1447).

*mcherry* sequence was amplified from pSJ9016 (Zuryn et al., 2014) using the primers F: 5’-CCCTTGGAGGGTACC-CAGCTG-3’ and R: 5’-CATTCCTCCCTTATACAATTCATCCATGCCACC-3’ and cloned into the NheI site of the subsequent plasmid.

*lin-15A cDNA* was amplified from genomic N2 DNA via PCR using the primers F5’- ATGTTG-GCTCCAGCGGCTCCAGCTAAAGATGTTGTCTCG-3’ and R 5’-CGCTTAAAAAATTGGCTCAGGCTTTGGATCC-3’. *lin-15A* cDNA was cloned by Gibson assembly using the primers F 5’-GGGAGGAATG-TTGGCTCCAGCGGCTCC-3’ and R 5’-CTGCGGCCGCTTAAAAAATTGGCTCAGGCTTTGG-3’.

For the *egl-5(6.5kb)p::lin-15A cDNA::SL2::mcherry* transgenic plasmid *lin-15A cDNA* was amplified from pSJ805 using the primers F: 5’-AAAAGATCTTATCTGACTCCGTGCATAC-3’ and R: 3’-AGGCCTTTATGTCCGGTGGCATCTC-5’and cloned into the Acc65I restriction site of pSJ962 Zuryn et al. (2014).

For the *egl-5(6.5kb)p::lin-15A cDNA dCterm::SL2::mcherry* transgenic plasmid *lin-15A cDNA* lacking the C-term was amplified from pSJ805 using the primers F: 5’-CTACGCCAGATCTGCTAGCGCG-3’ and R: 3’-AGGCCTTTATGTCCGGTGGCATCTC-5’ and cloned into the Acc65I restriction site of pSJ962.

For the *egl-5(6.5kb)p::lin-15A cDNA dNterm::SL2::mcherry* transgenic plasmid *lin-15A cDNA* lacking the N-term was amplified from pSJ805 using the primers F: 5’-AAAAGATCTTATCTGACTCCGTGCATAC-3’ and R: 3’-GAATTCATGCATAGGCCTGCG-5’ and cloned into the Acc65I restriction site of pSJ962.

For the *egl-5(6.5kb)p::lin-15A cDNA dZnF::SL2::mcherry* transgenic plasmid point mutations were induced at C212S and C215S using mutagenesis PCR with the primers F: 5’CCTATCTGACTCCGGGCATACTCTGC -3’ and R: 3’- GCAGAGTATGCCCGGAGTCAGATAGG -5’ in pSJ805. The mutated lin-15A cDNA was amplified using the primers F: 5’-AAAAGATCTTATCTG-ACTCCGTGCATAC-3’ and R: 3’-AGGCCTTTATGTCCGGTGGCATCTC-5’ and cloned into the Acc65I restriction site of pSJ962.

The *hsp-16.2p::lin-15A cDNA::mCherry* transgenic plasmid pSJ830 was generated by cloning the *mcherry::lin-15A cDNA* sequence in between the BstBI and PvuI sites of pSJ9016 Zuryn et al. (2014) into the AvaI site of pSJ805. *mcherry::lin-15A* cDNA was amplified from pSJ805 using the following primers 5’- CCGGGATTGGCCAAAGGACCCAAAGGTATGTTTCGAATG-3’ and 5’-CGGTCCTCCG-ATCGTTGTCAGAAGTAAGTTGGC-3’.

### *lin-35* tissue-specific rescue constructs

For the *egl-5(1.3kb)p::lin-35 cDNA::mkate* transgenic plasmid *pSJ835 lin-35 cDNA* was cloned by Gibson assembly into pSJ834 in between the *delta-pes-10* and *mkate* sequences using the primers F: 5’-ATGCCGAAACGAGCAGCCGATG-3’ and R: 5’-ATCTCCAGACCGTTCCATAGCG-3’.

For the *ugt-22p::lin-35 cDNA::mkate* transgenic plasmid 830bp of *ugt-22* promoter sequence was cloned into pSJ835 restriction sites HindIII and XbaI using the primers F: 5’AGCTTGCATGCC-TGCAGGTCGAC-3’ and R: 5’-CCGGTACCCTCCAAGGGTCCTC-3’.

For the *eft-3p::lin-35 cDNA::mkate* transgenic plasmid 610bp of *eft-3* promoter sequence was cloned into pSJ835 restriction sites HindIII and XbaI using the primers F: 5’- ATGAAATA-AGCTTGCATGCCTGCAGGCCTTGGTCGACGCACCTTTGGTCTTTTATTG-3’ and R: 5’-GGATCCTCTAGAGAGCAAAGTGTTTCCCAACT-3’.

For the *rab-3p::lin-35 cDNA::mkate* transgenic plasmid 1211bp of *rab-3* promoter sequence was cloned into pSJ835 restriction sites HindIII and XbaI using the primers F: 5’ ATGGATACGCTAACAACTTGGAAATGAAATAAGCTTATCTTCAGATGGGAGCAGTGGAC-3’ and R: 5’-CTACAGTAGCCCTATTTTCAGATGATCTCTAGAGGATCCCCGG-GGATTGGCCAAAGGACC–3”

For the *elt-2p::lin-35 cDNA::mkate* transgenic plasmid 2896bp of *elt-2* promoter sequence was cloned into pSJ835 restriction sites HindIII and XbaI using the primers F: 5’ GAAATAAGCTTTTGCATGCCTTCGTGTGATGG - 3’ and R: 5’- GGATCCTCTAGA-GATCTATAATCTATTTTCTAG – 3”

### Immunofluorescence

Whole-mount immunofluorescence was performed on synchronized L3 worms using a slightly modified Finney-Ruvkun procedure (Richard et al., 2011). Primary antibody mouse anti-EGL-5 (1/100) was a gift from Scott Emmons and rabbit anti-AJM-1 (1/400) was obtained from Developmental Studies Hybridoma Bank. Secondary staining was performed using either Cy3-conjugated (Jackson ImmunoResearch Laboratories) or Alexa 488-conjugated (Invitrogen) antibodies. Imaging was performed using a Yokogawa CSU-W1 Leica DMi8 spinning disk microscope.

### Western Blots

Whole worm protein extracts (mixed embryos and L1 larvae) prepared using RETSCH Mixer Mill MM 400) were resuspended into the double distilled water and mixed with appropriate volume of 4X-SDS loading buffer (Total- 50-100 l). Samples were heated at 95°C for 5 mins on a PCR machine followed by a quick centrifugation before loading on an SDS-PAGE. The SDS-PAGE (10-12%) was used to separate proteins. Samples were then transferred onto a PVDF membrane using standard wet western blot transfer methods. Blot transfer was achieved by running blot set-up at 120V for 120 at 4°C. This condition was sufficient for transfer for all molecular weight protein. After transfer, the membrane was blocked with TBST buffer with 5% skimmed milk with gentle shaking using rockers for 1hr at RT. The membrane was incubated with the appropriate primary antibody – Anti-LIN-15A Q2015 (1:1000 dilution in TBST buffer with 2.5% skimmed milk) and kept with gentle shaking using rockers for overnight at 4°C. The next day, the membrane was washed thrice at RT for 10 minutes in TBST buffer then incubated with HRP conjugated Goat anti-mouse secondary antibodies (GAM, made in-house (0.5 l in 10 ml of TBST (1/20000 dilution) and 5 ml used for one blot) for 120 mins at RT on mild shaking. Blots were washed thrice for 10 minutes in TBST buffer and developed by Thermo Scientific™ SuperSignal™ West Pico PLUS Chemiluminescent Substrate (Catalog number 34577) as manufacturer’s protocol.

### LIN-15A streptavidine and FLAG pull-downs

For both experiments, 1 ug of LIN15A-FLAG tagged purified protein was incubated with 400 ng of Biotinylated DNA or 400ng of Biotinylated Recombinant or 400ng of Biotinylated Native Mononucleosomes (from HeLa cells) in 100mM NaCl. Samples were pull down with either Streptavidine-agarose bead or FLAG-agarose beads resuspended in SB2X, loaded on a 8%-12% SDS PAGE gradient gels and revealed with anti-FLAG or anti-H3 antibodies. For the FLAG pull-down, half of the pulled-down sample was spotted on a nylon membrane, UV crosslinked and revealed with an anti-streptavidine-HRP conjugated antibody.

### Generation of *lin-15A*(*fp31*) and *lin-15A*(*syb978*) alleles

Constructs containing Cas9 and sgRNAs with PAM sites before *lin-15A* start codon and after *lin-15* TATA box were injected together with Co-CRISPR sgRNA (pML3441/3442 against dpy-10) and repair template (OCG250 *rol-6*), as well as co-injection marker pCF390 into *lin-65(n3411); syIs63*. In F1 worms were selected for [Dpy]/[Rol] and co-injection marker, in F2 worms were selected for [Muv] phenotype and validated by sequencing. *lin-65(n3411)*, Co-CRISPR constructs and co-injection marker were crossed out or negatively selected to obtain *lin-15A(fp31); syIs63*.

The following constructs containing the sgRNAs were used:

- pSJ407 sgRNA 1.6 *lin-15A* start
- PSJ408 sgRNA 1.9 *lin-15A* start
- pSJ409 sgRNA 2.6 *lin-15A* start
- pSJ410 sgRNA 2.8 *lin-15A* start
- pSJ411 sgRNA 3.7 *lin-15A* 3’UTR
- pSJ412 sgRNA 3.8 *lin-15A* 3’UTR
- pSJ413 sgRNA 4.8 *lin-15A* stop
- pSJ414 sgRNA 4.8 *lin-15A* stop

Generation of endogenously tagged LIN-15A::mCherry *syb978* was outsourced to Suny Biotech. The *mCherry* sequence was added in Nter, right after *lin-15A* ATG, and separated by the ‘6xHis’ plus ‘linker’ sequence CATCACCATCACCATCACGGAGGAAACGGAAA-CAACGCTGCCGGA

### Cell fate challenge assays

Blastomere multipotency was assessed through the overexpression of the muscle-determining transcription factor HLH-1 under a heat-shock inducible promoter following Fukushige and Krause’s (Fukushige and Krause, 2005) procedure. Strains carried an integrated array containing *hlh-1* cDNA under the control of the *hsp-16.41* promoter (Stringham et al., 1992) and a *rol-6* phenotypic marker as well as a muscle-specific fluorescent reporter *ccIs4251*, inducing the expression of the GFP in the nucleus and the mitochondria of muscle-determined blastomeres. The precise protocol for this assay can be found in (Lambert and Jarriault, *in prep*).

### Chromatin-Immuno Precipitation sequencing (ChIP-seq) of LIN-15A, LIN-15B and LIN-35

ChIP-seq were performed in duplicates at minima. Synchronization of the L1 larvae was obtained through series of hatch-pulses. Briefly, WT, *lin-15A(fp31)*, *lin-15B(n744)* and *lin-35(n745)* animals were grown on peptone plates with *E. coli* HB101 as a food source (http://www.wormbook.org/chapters/wwwtrans-formationmicroinjection/transformationmicroinjection.htmld0e878) and then bleached. Eggs were deposited on empty NGM plates which were washed with M9 every three hours. Collected animals were immediately frozen into liquid nitrogen (‘pearls’). Samples from the successive washes were pulled together. *fp31* is a deletion of the *lin-15A* loci (*lin-15B* unchanged). *n744* and *n745* are premature stop codons, respectively in LIN-15B and LIN-35. Worm pearls were ground into a powder using a RETSCH Mixer Mill MM 400 (13 seconds at 25 Htz) and subsequently pre-incubated in PBS. Fixation was performed by addition of 1.5 mM EGS (Pierce 21565) for 4 minutes, followed by the addition of formaldehyde to a final concentration of 1%, and incubated for a further 4 minutes. Fixation was stopped by the addition of 0.125 M glycine for 5 minutes. Fixed tissues were washed 2X with PBS with protease inhibitors (Roche EDTA-free protease inhibitor cocktail tablets 05056489001) and once in FA buffer (50 mM HEPES pH7.5, 1 mM EDTA, 1% Triton X-100, 0.1% sodium deoxycholate, and 150 mM NaCl) with protease inhibitors (FA+), then resuspended in 1 mL FA+ buffer per 1 mL of ground worm powder. The extracts were sonicated to an average size of 250 base pairs using a Bioruptor Pico (Diagenode), and 15 µg of DNA was used per ChIP reaction. The following antibodies (all from the Ahringer lab) were used for the Chromatin-Immunoprecipitation: Q2015 – anti-LIN-15A (this study), Q2330 – anti-LIN-15B (Rechtsteiner et al., 2019), Q2001 – anti-LIN-35 (Gal et al., 2021). ChIP-seq and library preparations were performed as described in (Jänes et al., 2018).

Genome and canonical gene annotations were downloaded from Wormbase (release 185), retaining only protein coding, pseudogenes and lincRNAs. ChIP-seq data were preprocessed using trim galore (available at https://github.com/FelixKrueger/TrimGalore, version 0.6.7) and mapped using bwa mem (Li and Durbin, 2009) (version 0.7.17). RPGC-normalized (reads per genomic content) coverage tracks from mapq10 reads were generated using bamCoverage from the deepTools2 suite (Ramirez et al., 2016)(version 3.5.5). Peaks from each replicate were called using MACS2 (Zhang et al., 2008) (version 2.2.9.1) using no control sample and the following parameters: –call-summits –bdg –SPMR –gsize ce –keep-dup all –nomodel. To define a set of peaks for each wild-type or mutant condition, MACS2 summits with a score >6 were first extended to 100bp on each side. Peaks from each replicate condition were then intersected using the intersect function from BED-tools (Quinlan and Hall, 2010) (version 2.30.0), and the intersected peaks retained only if the MACS2 score from each replicate was between 0.1- and 10-fold of the score from the other replicate (or the average of the other replicates, when three replicates were available). This was done to mitigate the impact of between-replicate variation in the quantification of differentially bound regions (see below). Intersected peaks were then rescaled to 200bp around their midpoint. PCA analysis and correlation plots among wild-type and mutant replicates from each factor were carried out using the deepTools2 suite.

Sites showing differential enrichment of each factor between the wild-type and each mutant were defined using the R (version 4.2.0) package DESeq2 (Love et al., 2014) (version 1.38.0). For this analysis, all peaks for each factor called in the wild type and mutant conditions were combined using the merge function from BEDtools, and the ChIP-seq data were quantified in each peak using summarizeOverlaps (mode=”Union”). Annotation of the promoters were performed according to (Serizay et al., 2020) for comparison with the LIN-15B and LIN-35 peaks in the WT backgrounds from (Gal et al., 2021), and using wormbase WBCel235 annotations to identify the closest predicted gene (intragenic, or upstream peaks ≤2kb from the ATG).

Gene Ontology (GO) and Kyoto Encyclopedia of Genes and Genomes (KEGG) pathway enrichment analyses were carried out with ShinyGO 0.85 (Ge et al., 2020; Luo and Brouwer, 2013; Kanehisa et al., 2021), using *C. elegans* genome as background. Pathways & GOs were selected based on FDR and sorted by their respective Fold Enrichment.

## Acknowledgements

Some strains were provided by the CGC, which is funded by NIH Office of Research Infrastructure Programs (P40 OD010440), and by the National BioResource Project, which is funded by the Japanese government. We thank Francesc Castro and Marta Gut for help with WGS of *C. elegans* mutants, and all members of the Jarriault team for critical discussions and constant support. Research was supported by the Institut National de la Santé et de la Recherche Médicale (PRI NeuroLicencing), and the European Research Council (ERC PlastiCell 648960) to SJ, and by the ANR-10-LABX-0030-INRT grant, a French State fund managed by the Agence Nationale de la Recherche under the Investissements d’Avenir program ANR-10-IDEX-0002-02, and SFRI-STRAT’US project (ANR-20-SFRI-0012) under the framework of the France 2030 Program. SFB. is a recipient of a La Ligue Contre le Cancer fellowship. MCM is a recipient of a Fondation pour la Recherche Médicale fellowship. JL is a recipient of a CDSN and an Association pour le Recherche sur le Cancer fellowships, an EMBO short exchange fellowship and a Company of Biologists travelling fellowship. SJ is a CNRS research director.

## CRediT

**Conceptualization :** SFB, MCM, JL, SJ; **Investigation :** SFB, MCM, JL, SKS, AA, SR, SHY, JMS, MP, RM, JA; **Methodology :** JL; **Resources :** SFB, MCM, JL, SHY, JMS, MP; **Formal analysis :** SFB, MCM, JL, FC, SLG; **Data curation :** JL, FC, SLG, SJ; **Supervision :** SFB, MCM, JL, SJ; **Visualization :** SFB, JL, SJ; **Writing – original draft :** SFB, JL, SJ; **Project administration :** SJ; **Funding acquisition :** SJ

## Data and resource availability

*On-going*

