## Supplementary material for "The SynMuvA *lin-15A* licenses natural transdifferentiation by antagonizing identity safeguarding mechanisms": SI figures & legends

**Figure S1: SynMuvAs LIN-15A and LIN-56 are required for Y-to-PDA initiation. A** Allele complementation shows that fp22 and n767 are recessive, null alleles of lin-15A. **B** LIN-15A and LIN-56 proteins form a dimer in vivo: a 115kDa LIN-15A band (yellow box) is observed rather than the 80kDa expected. Likewise, when using the fusion protein LIN-15A::mCherry, a 135 to 140kDa (red box) band is observed instead of a 110kDa band. As this band is approximatevely 30kDa higher than the lin-15A alone band, and as 28kDa is the expected weight for mCherry, suggest that these bands that are higher than expected do correspond to the LIN-15A::mCherry fusion. lin-56 is 37kDa, which corresponds to the added weight in these bands, suggesting the what is detected on these WB is a very stable LIN-15A-LIN-56 complex. Q2015 targets the LIN-15A / LIN-56 dimer as the same aspecific bands are found in the two corresponding mutants. **C** *lin-15A* and *lin-56* are the only SynMuvAs involved in Y-to-PDA transdifferentiation. Inactivation via mutant alleles or RNA interference of the other SynMuvAs does not induce any defect, nor severe enhance the Y-to-PDA defective phenotypes of *lin-15A(n767)* mutants. X2 and Fisher exact tests with Bonferroni corrections for proportion comparisons. **D** Representative pictures of the rectal area during the L1 stage. The rectal slit is indicated by the red bar. Rectal-epithelial markers *col-34* and *lin-26* expression is not affected by loss of *lin-15A* prior to Td initiation. Scalebars = 5 µm. **E** Expression of the transgenes used in LIN-15A rescue-experiments are expressed in Y prior to Td initiation. Images of the rectal area in L1. Scalebar = 5 µm. **F** LIN-15A is nuclear and ubiquitously / widely expressed during later embryonic stages / early larval development. LIN-15A is expressed in Y during the L1 stage, prior to Td. The rectal slit is indicated in red. Scalebars embryo = 20 µm. Scalebars L1 larvae = 10 µm.

**Figure S2: LIN-15A does not regulate CEH-6 nor SOX-2 expression. A** CEH-6 expression is not affected by the loss of *lin-15A* (n WT=45 / n n767=50). **B** SOX-2 expression is not affected by the loss of *lin-15A* (n WT=48 / n n767=54).

**Figure S3: LIN-15A function in Y-to-PDA is independant of LIN-3. A** Loss of function (*mt378* and *e1417*) and gain of function (*n4441*) alleles of *lin-3* do not induce defects in Y-to-PDA transdifferentiation. **B** LIN-15A function in Y-to-PDA is independent of *lin-3*. The penetrance of *lin-15A(n767)* is not affected by loss or gain of function mutations of *lin-3*. X2 and Fisher exact tests for proportion comparisons.

**Figure S4: Title to be determined. A** Loss of the SynMuvBs LIN-15B, LIN-35 or LIN-13 suppresses the defects of *lin-56(n2728)* mutants. X2 and Fisher exact tests with Bonferroni corrections for proportion comparisons. **B i** Tissue-specific rescue of LIN-35/pRb. Ubiquitous and rectal-specific rescues of LIN-35/pRb strongly anti-suppress Td defects in double *lin-35(n745); lin-15A(n767)* mutants. Intestinal and neuronal rescues have weaker and/or inconsistent effect. **ii** Expression of the different constructs during the L1 stage (prior to Td initiation) used in the *lin-35* rescue experiments in the rectal area of the animals. Scalebar = 10 µm. **C i** Expression levels in Y of endogenously-tagged LIN-35/pRb in L1 (prior to Td) are unchanged in *lin-15A(n767)* compared to control animals suggesting a post-translational regulation. Shapiro–Wilk test for normality. Student t-test for mean comparisons. **ii** Representative images of the rectal area in transmission, and fluorescence: in red the rectal marker EGL-5::mCherry in green the endogenous LIN-35::GFP. The Y cell is indicated as well as P12.pa which replaces Y in the rectum after Td. P12.pa does not express EGL-5::mCherry. The rectal slit is indicated with the yellowbar. Scalebar = 10 µm. **D i** The expression levels in Y of endogenously-tagged MET-2 in L1 (prior to Td) are slightly decreased in *lin-15A(n767)* compared to control animals demonstrating that LIN-15A does not repress MET-2 expression. Student t-test for mean comparisons. Representative images of the rectal area in transmission, and fluorescence: in green the Y marker HLH-16::GFP – in red the endogenous MET-2::mCherry. The Y cell is indicated in white, the rectal slit in yellow. Scalebar = 10 µm.

**Figure S5: The two NuRD complexes of *C. elegans* have distinct an opposite effects on the plasticity of Y. A.** Overview of the components of the two NuRD and of the MEC complexes and their respective influence on cellular plasticity. **B** Loss of the Mi2 homolog CHD-3, specific to one of the two NuRDs, enhances the Td defects in *lin-15A(n767)* mutants but has no effect in the WT background. The *chd-3(eh4)* allele, which contains a 2 kb deletion that removes most of the CHD-3 helicase domain, is likely null (Käser-Pebernard 2016). Fisher exact test for proportion comparisons. **C** Loss of the CHD-3 also enhances the Td defects in *egl-27(ok151)* mutants. Fisher exact test for proportion comparisons. **D** CHD-3 acts as an enhancer for cellular plasticity in the two parallel pathways.

**Figure S6: LIN-15A binds chromatin. A** Streptavidine pull-down experiment with 1 ug of LIN15A-FLAG tagged purified protein incubated with 400 ng of Biotinylated DNA (DNA-Bio) or 400ng of Biotinylated Recombinant (Rec Mononucl-Bio) or 400ng of Biotinylated Native Mononucleosomes and pulled down with Streptavidine agarose beads. Loaded on a 8%->12% SDS PAGE gradient gel and revealed with anti-FLAG or anti-H3 antibodies. **B** FLAG pull down experiment with 1 ug of LIN15A-FLAG tagged purified protein incubated with 400 ng of Biotinylated DNA (DNA-Bio) or 400ng of Biotinylated Recombinant (Rec Mononucl-Bio) or 400ng of Biotinylated Native Mononucleosomes and FLAG agarose beads. ½ of the pull down was revealed with anti-FLAG or anti-H3 antibodies. ½ of the pulled down was revealed with an anti-streptavidine-HRP coniugated antibody. **C** LIN-15A binds mainly onto gene promoters (59%), similarly to LIN-15B (53,7%) and LIN-35 (56,6%). 5382/4211 (LIN-35), 6540/5172 (LIN-15B), 5295/4508 (LIN-15A) total peaks/genes were bound. **D** Overlap of the genes bound by LIN-15A, LIN-15B and LIN-35/pRb. All three factors overlap on a large fraction of their bound genes. **E** Biological Processes (**i**) and Cellular Components (**ii**) GO enrichment of the 2770 genes bound by the three factors, as obtained with

ShinyGO 0.85. BP GOs are enriched in terms associated with biosynthesis/biogenesis, cellular organisation and regulation of gene expression. CC GOs are enriched in terms associated with the nucleus/chromosomes and membranes - as well as in transport, metabolic (mitochondria, autophagy, mTOR) and membrane-related KEGGs (Table S2).

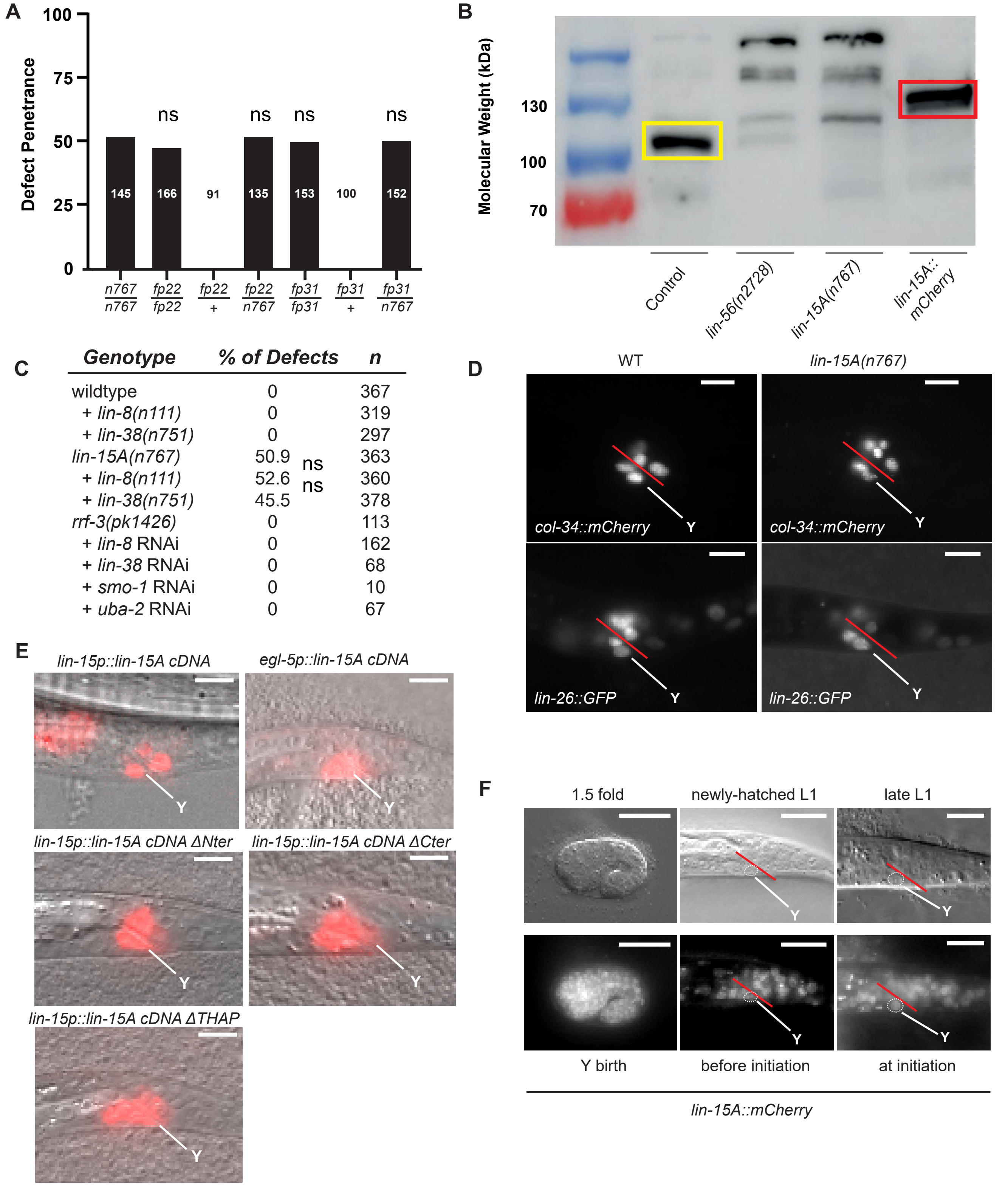


| Genotype | % of Defects | | n |
|---|---|---|---|
| wildtype | 0 | | 367 |
| + *lin-8(n111)* | 0 | | 319 |
| + *lin-38(n751)* | 0 | | 297 |
| *lin-15A(n767)* | 50.9 | | 363 |
| + *lin-8(n111)* | 52.6 | ns | 360 |
| + *lin-38(n751)* | 45.5 | ns | 378 |
| *rrf-3(pk1426)* | 0 | | 113 |
| + *lin-8* RNAi | 0 | | 162 |
| + *lin-38* RNAi | 0 | | 68 |
| + *smo-1* RNAi | 0 | | 10 |
| + *uba-2* RNAi | 0 | | 67 |

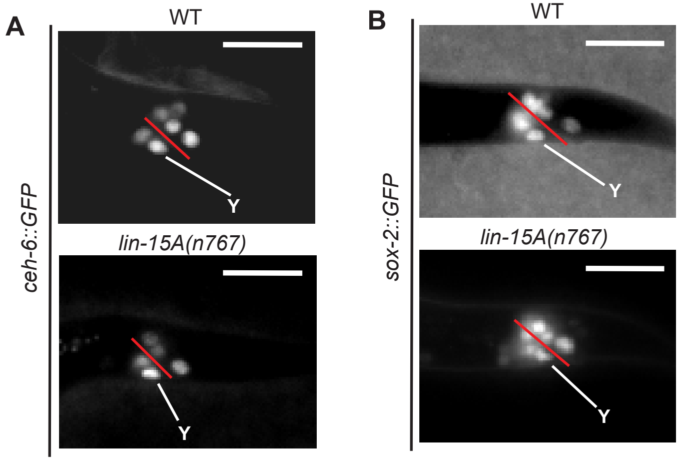

A
WT
Y
lin-15A(n767)
Y
ceh-6::GFP
B
WT
Y
lin-15A(n767)
Y
sox-2::GFP

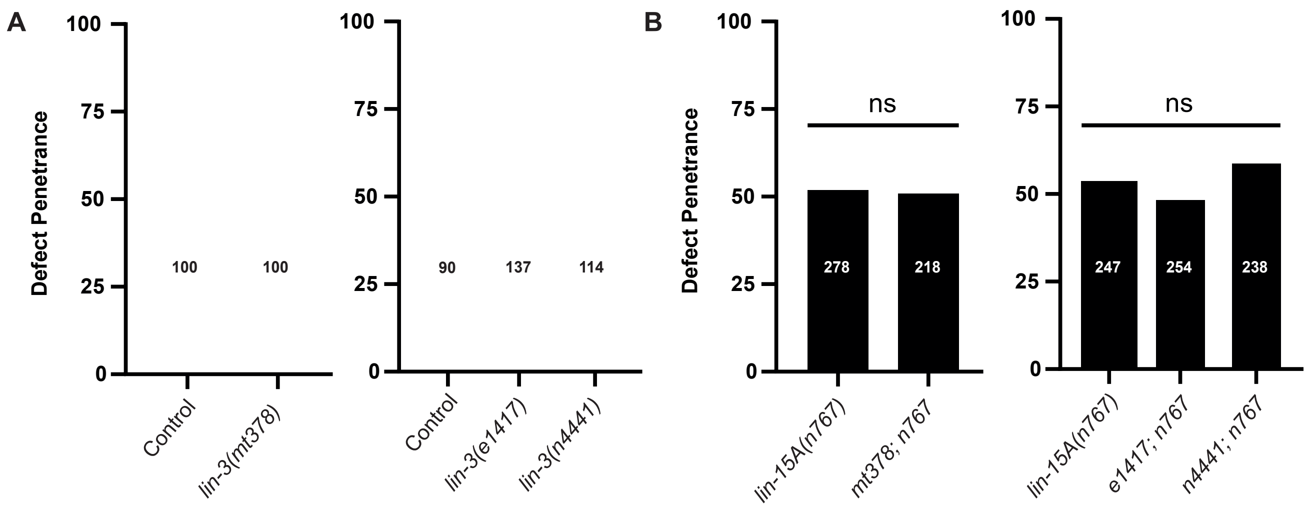
A
B
Defect Penetrance
100
75
50
25
0
Control
lin-3(mt378)
100
100
lin-3(e1417)
lin-3(n4441)
90
137
114
ns
lin-15A(n767)
mt378; n767
278
218
e1417; n767
n4441; n767
247
254
238

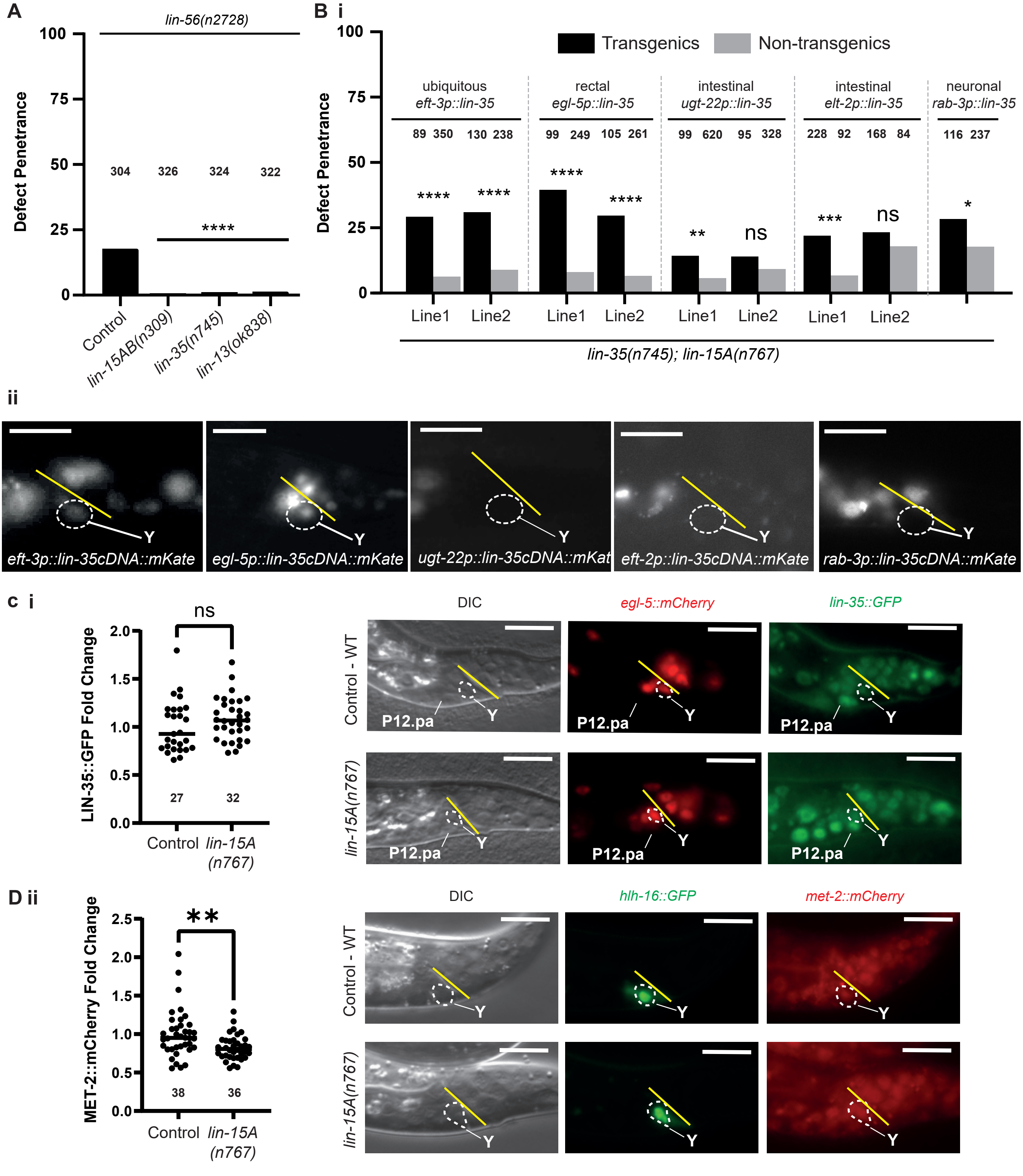
A
lin-56(n2728)
Defect Penetrance
304 326 324 322
****
Control
lin-15AB(n309)
lin-35(n745)
lin-13(ok838)
B i
Transgenics
Non-transgenics
ubiquitous
eft-3p::lin-35
rectal
egl-5p::lin-35
intestinal
ugt-22p::lin-35
intestinal
elt-2p::lin-35
neuronal
rab-3p::lin-35
89 350 130 238 99 249 105 261 99 620 95 328 228 92 168 84 116 237
Line1 Line2 Line1 Line2 Line1 Line2 Line1 Line2
lin-35(n745); lin-15A(n767)
ii
eft-3p::lin-35cDNA::mKate
egl-5p::lin-35cDNA::mKate
ugt-22p::lin-35cDNA::mKat
eft-2p::lin-35cDNA::mKate
rab-3p::lin-35cDNA::mKate
c i
LIN-35::GFP Fold Change
ns
27 32
Control
lin-15A (n767)
DIC
egl-5::mCherry
lin-35::GFP
Control - WT
lin-15A(n767)
P12.pa
Y
D ii
MET-2::mCherry Fold Change
**
38 36
Control
lin-15A (n767)
DIC
hlh-16::GFP
met-2::mCherry
Control - WT
lin-15A(n767)
Y

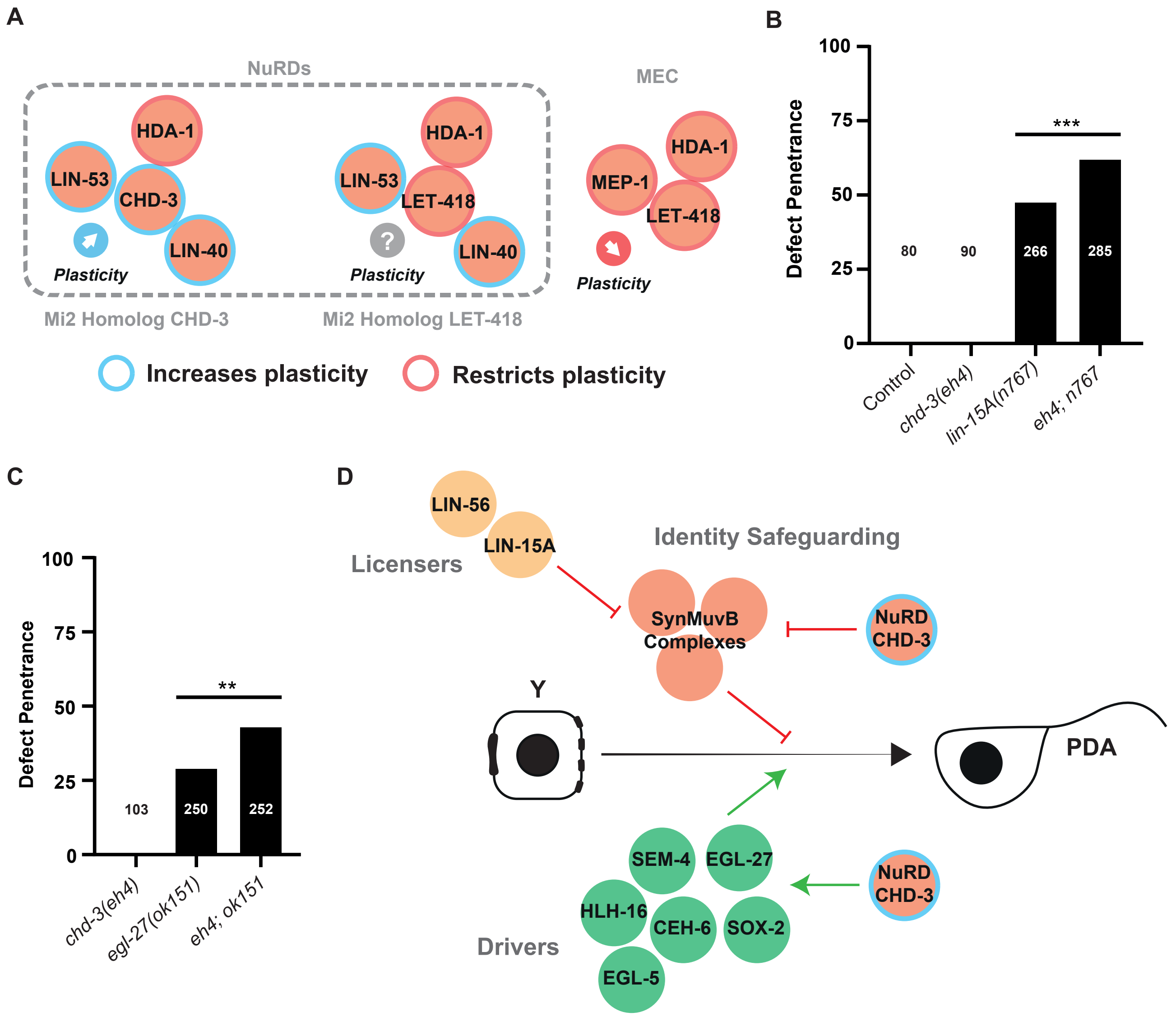
A
NuRDs
HDA-1
LIN-53
CHD-3
LIN-40
Plasticity
Mi2 Homolog CHD-3
HDA-1
LIN-53
LET-418
LIN-40
?
Plasticity
Mi2 Homolog LET-418
MEC
HDA-1
MEP-1
LET-418
Plasticity
Increases plasticity
Restricts plasticity
B
Defect Penetrance
100
75
50
25
0
***
80
90
266
285
Control
chd-3(eh4)
lin-15A(n767)
eh4; n767
C
Defect Penetrance
100
75
50
25
0
**
103
250
252
chd-3(eh4)
egl-27(ok151)
eh4; ok151
D
LIN-56
LIN-15A
Licensers
Identity Safeguarding
SynMuvB Complexes
NuRD CHD-3
Y
PDA
SEM-4
EGL-27
HLH-16
CEH-6
SOX-2
EGL-5
Drivers
NuRD CHD-3

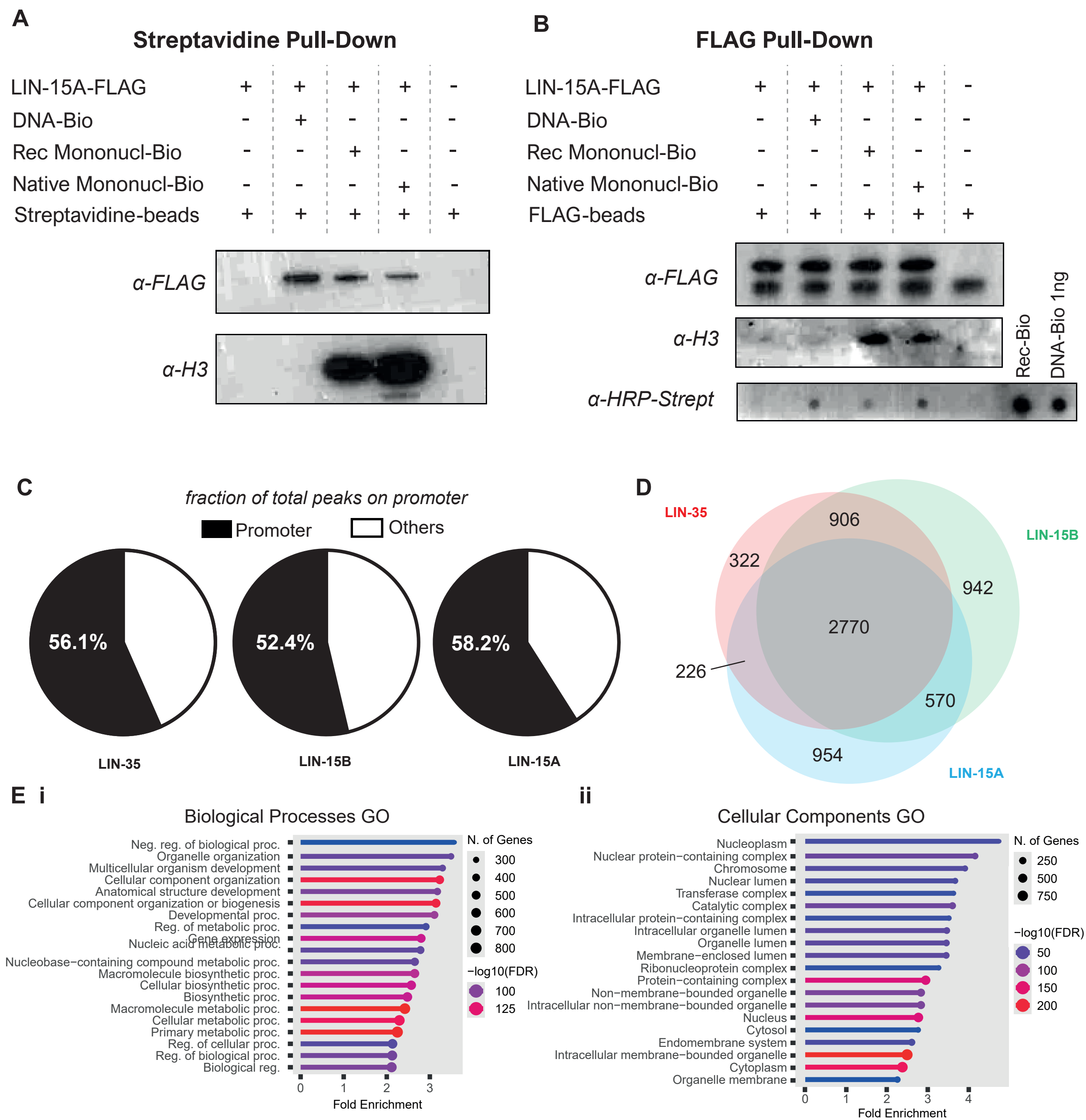
A
Streptavidine Pull-Down
LIN-15A-FLAG + + + + -
DNA-Bio - + - - -
Rec Mononucl-Bio - - + - -
Native Mononucl-Bio - - - + -
Streptavidine-beads + + + + +
α-FLAG
α-H3
B
FLAG Pull-Down
LIN-15A-FLAG + + + + -
DNA-Bio - + - - -
Rec Mononucl-Bio - - + - -
Native Mononucl-Bio - - - + -
FLAG-beads + + + + +
α-FLAG
α-H3
α-HRP-Strept
Rec-Bio
DNA-Bio 1ng
C
fraction of total peaks on promoter
Promoter
Others
56.1%
52.4%
58.2%
LIN-35
LIN-15B
LIN-15A
D
LIN-35
LIN-15B
LIN-15A
906
322
942
2770
226
570
954
E i
Biological Processes GO
Neg. reg. of biological proc.
Organelle organization
Multicellular organism development
Cellular component organization
Anatomical structure development
Cellular component organization or biogenesis
Developmental proc.
Reg. of metabolic proc.
Gene expression
Nucleic acid metabolic proc.
Nucleobase−containing compound metabolic proc.
Macromolecule biosynthetic proc.
Cellular biosynthetic proc.
Biosynthetic proc.
Macromolecule metabolic proc.
Cellular metabolic proc.
Primary metabolic proc.
Reg. of cellular proc.
Reg. of biological proc.
Biological reg.
0 1 2 3
Fold Enrichment
N. of Genes
300
400
500
600
700
800
−log10(FDR)
100
125
ii
Cellular Components GO
Nucleoplasm
Nuclear protein−containing complex
Chromosome
Nuclear lumen
Transferase complex
Catalytic complex
Intracellular protein−containing complex
Intracellular organelle lumen
Organelle lumen
Membrane−enclosed lumen
Ribonucleoprotein complex
Protein−containing complex
Non−membrane−bounded organelle
Intracellular non−membrane−bounded organelle
Nucleus
Cytosol
Endomembrane system
Intracellular membrane−bounded organelle
Cytoplasm
Organelle membrane
0 1 2 3 4
Fold Enrichment
N. of Genes
250
500
750
−log10(FDR)
50
100
150
200

| Factor | Complex |
|---|---|
| LIN-35 | DREAM |
| DPL-1 | DREAM |
| LIN-9 | DREAM |
| LIN-37 | DREAM |
| LIN-54 | DREAM |
| LIN-52 | DREAM |

| Factor | Complex |
|---|---|
| EFL-1 | DREAM |
| LIN-53 | DREAM, NuRD and others |
| LIN-15B | DREAM associated |
| LIN-36 | DREAM associated |
| LET-418 | NuRD, MEC |
| HDA-1 | NuRD, MEC |

| Factor | Complex |
|---|---|
| MEP-1 | MEC |
| HPL-2 | Heterochromatin |
| MET-2 | Heterochromatin |
| LIN-61 | Heterochromatin |
| LIN-13 | Heterochromatin |
| LIN-65 | Heterochromatin |
